# A cell atlas of the fly kidney

**DOI:** 10.1101/2021.09.03.458871

**Authors:** Jun Xu, Yifang Liu, Hongjie Li, Alexander J. Tarashansky, Colin H. Kalicki, Ruei-Jiun Hung, Yanhui Hu, Aram Comjean, Sai Saroja Kolluru, Bo Wang, Stephen R Quake, Liqun Luo, Andrew P. McMahon, Julian A.T. Dow, Norbert Perrimon

## Abstract

Like humans, insects rely on precise regulation of their internal environments to survive. The insect renal system consists of Malpighian tubules and nephrocytes that share similarities to the mammalian kidney. Studies of the *Drosophila* Malpighian tubules and nephrocytes have provided many insights into our understanding of the excretion of waste products, stem cell regeneration, protein reabsorption, and as human kidney disease models. Here, we analyzed single-nucleus RNA sequencing (snRNA-seq) data sets to characterize the cell types of the adult fly kidney. We identified 11 distinct clusters representing renal stem cells (RSCs), stellate cells (SCs), regionally specific principal cells (PCs), garland nephrocyte cells (GCs) and pericardial nephrocytes (PNs). Analyses of these clusters revealed many new interesting features. For example, we found a new, previously unrecognized cell cluster: lower segment PCs that express *Esyt2*. In addition, we find that the SC marker genes *RhoGEF64c*, *Frq2*, *Prip* and *CG10939* regulate their unusual cell shape. Further, we identified transcription factors specific to each cluster and built a network of signaling pathways that are potentially involved in mediating cell-cell communication between Malpighian tubule cell types. Finally, cross-species analysis allowed us to match the fly kidney cell types to mouse kidney cell types and planarian protonephridia - knowledge that will help the generation of kidney disease models. To visualize this dataset, we provide a web-based resource for gene expression in single cells (https://www.flyrnai.org/scRNA/kidney/). Altogether, our study provides a comprehensive resource for addressing gene function in the fly kidney and future disease studies.

## INTRODUCTION

The functions of excretory systems are to remove toxins from the body and maintain homeostatic balance. For example, mammalian kidneys play important roles in several physiological processes, including maintaining water fluid homeostasis, removing metabolic waste products, controlling blood pressure, regulating blood cell composition, and regulating bone mineralization (Nielsen et al., 2012). Although the excretory systems of various animals have differences, they typically have in common two activities: filtration and tubular secretion/reabsorption (Denholm and Skaer, 2009). In mammals, the mature kidney consists of two connected parts: a nephron, derived from the metanephric mesoderm, and a collecting tubule derived from the ureteric bud (Nielsen et al., 2012; McMahon, 2016). The *Drosophila* renal system is composed of separated filtration nephrocytes and Malpighian (renal) tubules (Miller et al., 2013). In *Drosophila*, about 25 garland cell nephrocytes (GCs) and 120 pericardial nephrocytes (PNs) are found at the end of embryogenesis and are maintained during development into the adult stage. GCs form a ring around the junction between the proventriculus and esophagus, whereas PNs are located on both side of the heart tube (Helmstädter et al., 2017). These two types of nephrocytes, although derived from different cell lineages, share morphological, functional, and molecular features with podocytes, which form the glomerular filter in vertebrates, and possess a protein sequestration activity reminiscent of the proximal tubule (Helmstädter et al., 2017). The adult Malpighian tubules, considered to be analogous to the renal tubular system, develop from the ectodermal hindgut primordium and visceral mesoderm and consist of two pairs of epithelial tubes that empty into the hindgut at its junction with the posterior midgut (Jung et al., 2005).

*Drosophila* Malpighian tubules and nephrocytes have been used to model human kidney diseases. Previously, a screen for genes involved in renal function identified over 70 genes required for nephrocyte function (Zhang et al., 2013). In addition, 30 human causative genes involved in Steroid resistant nephrotic syndrome (SRNS) have been analyzed in fly nephrocytes. Among them, *Cubilin* (*Cubn*) was found to be required for nephrocytes endocytosis (Hermle et al., 2017). Further, the coenzyme Q10 (CoQ10) biosynthesis gene *Coq2*, involved in regulating the morphology of slit diaphragm, and ROS formation, contribute to a pathomechanism of COQ2-nephropathy (Hermle et al., 2017). In addition to model numerous human renal conditions such as chronic kidney disease and kidney stones, the Malpighian tubule is also an excellent model in which to study the neuroendocrine control of renal function and rapid fluid transport (Cohen et al., 2020).

Single-nucleus (snRNA-seq) and single-cell (scRNA-seq) RNA sequencing give us an opportunity to understand the cellular make-up of many organ systems, including the kidney. The mammalian kidney is composed of cell types with unique functions. Podocytes regulate the passage of proteins, and the function of principal cells and intercalated cells in the collecting duct balance systemic water, pH, and salt levels (Garg, 2018; Pearce et al., 2015; Roy et al., 2015). A detailed scRNA-seq study defined the whole landscape of the mouse kidney, with 32 distinct clusters of ontology (Ransick et al., 2019). Finally, scRNA-seq data can be used for the analysis of pseudotemporal ordering of cells, which can provide information about developmental trajectories of cellular lineages.

Here, we performed snRNA-seq of the adult *Drosophila* male and female kidney system to characterize the organization, origins and diversity of the various cell types. Specifically, we identified 11 distinct clusters representing renal stem cells (RSCs), stellate cells (SCs), principal cells (PCs), garland nephrocytes cells (GCs) and pericardial nephrocytes (PNs), and provide gene expression level data at single cell resolution. In addition, based on the snRNA-seq data, we identified a set of genes involved in regulating cell shape of SCs. We also analyzed cell-to-cell communication between clusters, cluster-specific transcription factors, and metabolic differences between clusters. Importantly, performing a cross-species analysis between the fly kidney, planarian protonephridia and mouse kidney allowed us to map kidney cell types across species. We also analyzed human kidney disease genes at the cluster level in the fly kidney. Finally, we built a web-based visualization resource (https://www.flyrnai.org/scRNA/kidney/) that allows users to browse snRNA-seq data and query the expression of genes of interest in different cell types.

## RESULTS

### snRNA-seq identifies 11 distinct clusters in the adult *Drosophila* kidney

The fly kidney consists of Malpighian tubules and nephrocytes that are located in different regions of the body. Malpighian tubules branch from two common ureters that drain into the gut at the midgut/hindgut junction. Nephrocytes represent garland nephrocytes cells (GCs) located near the esophagus and proventriculus, and pericardial nephrocytes (PNs) located in the abdominal tissue (Fig. 1A). As part of the Fly Cell Atlas (FCA) project, we dissected male and female Malpighian tubules (Li et al., 2021) and annotated the cell types. In addition, as nephrocytes were not included in the FCA, we performed snRNA-seq of both GCs and PNs. To visualize GCs and PNs during dissection, we expressed *UAS-GFP.nls* under the control of *Dot-Gal4,* which is expressed in both cell types. In total, 150 male and 150 female tissues were dissected. Subsequently, four independent samples were processed for single nucleus isolation and the mRNAs were barcoded and sequenced (Fig. 1A). We successfully recovered 12,166 cells in the tubules. We also identified nephrocyte cell clusters that include a GC cluster with 41 nuclei and a PN cluster with 93 nuclei. Details on the number of cells and statistics are summarized in Supplementary Table 1.

**Figure 1.**
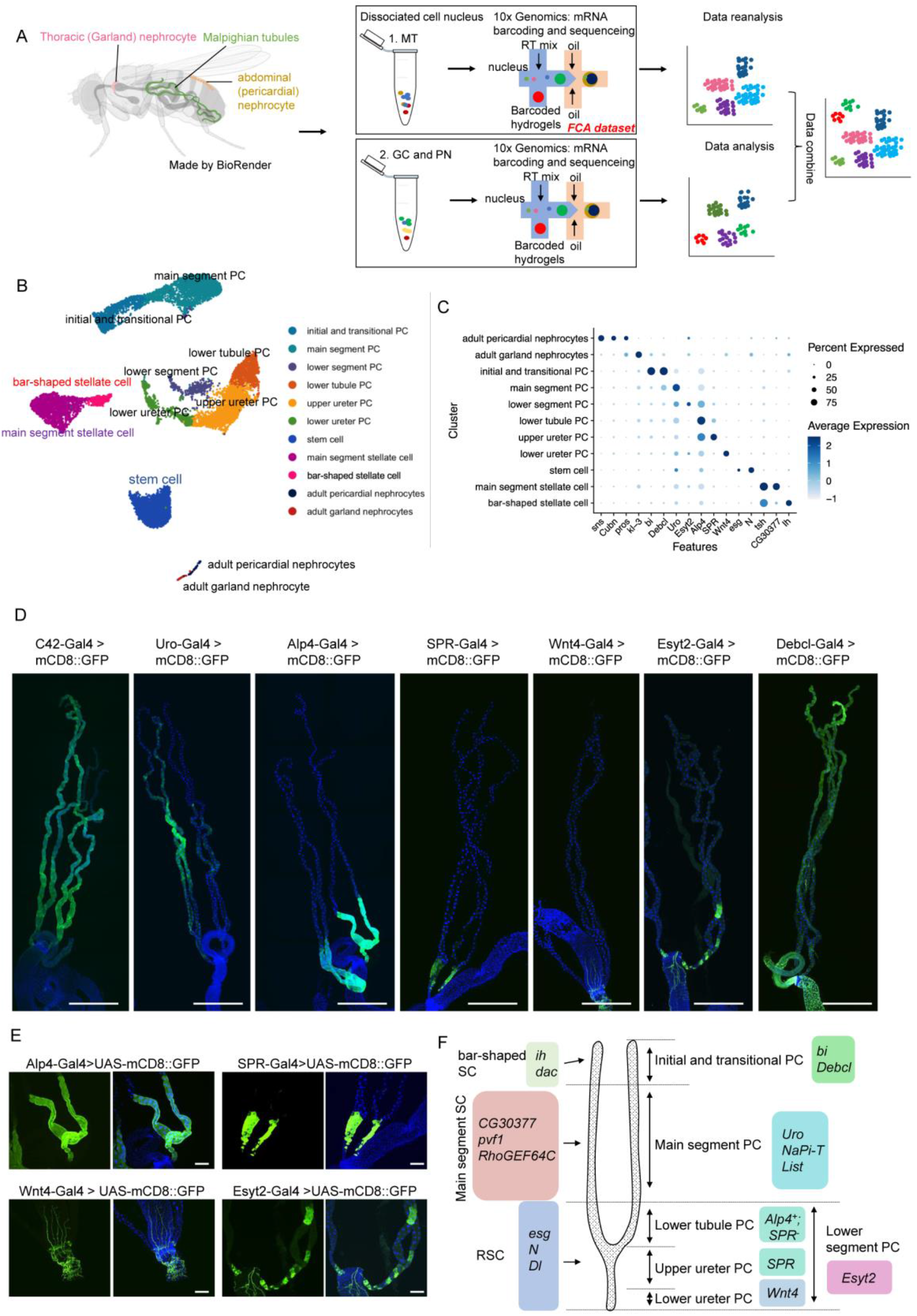
snRNA-seq analysis of the fly adult kidney. (A) Experimental design. The location of the Malpighian tubules and two types of nephrocytes are shown. Nuclei from Malpighian tubules and nephrocytes were processed separately and encapsulated using 10x Genomics. Data analysis was conducted independently and then combined to generate a single UMAP of the “fly kidney”. Note, however, that nephrocytes and tubules are not physically associated in vivo. (B) 11 distinct cell clusters were annotated on the UMAP. (C) Expression levels and percentage of cells expressing the marker genes in each cluster are shown as a dot plot. (D) GFP expression under the control of Gal4 lines specific for each of the six PC clusters. C42-Gal4 is expressed in all PCs. Scale bars = 500 μm. (E) Zoom-in of D panels to show the local features. Scale bars = 100 μm. (F) Malpighian tubule cell types are identified based on differentially expressed marker genes.

We identified 11 distinct clusters representing renal stem cells (RSCs), stellate cells (SCs), regionally specific principal cells (PCs), and nephrocyte cells (GCs and PNs) (marker genes listed in Supplementary Table 2). Note that the tubule snRNA-seq data were independently annotated at Harvard and FCA with highly concordant results (Fig. S1). We annotated the clusters based on previous knowledge and validation of novel marker genes (Supplementary Table 3). Previous studies have characterized enhancer trap lines that identified six distinct regions and genetically separable cell types in the tubules (Sözen et al., 1997). In addition, we validated new markers, identified as cluster-specific, by driving fluorescent reporters with the appropriate GAL4 lines (Supplementary Table 3).

The Malpighian tubule stem cell cluster is defined by the expression of *escargot* (*esg*), *Notch* (*N*) and *Delta* (*Dl*) genes (Wang and Spradling, 2020). The two stellate cell clusters (main segment SCs and bar-shaped SCs) both express *teashirt* (*tsh*), *kinin receptor (lkr)* and *Secretory chloride channel* (*SecCl*) (Denholm et al., 2013; Radford et al., 2002; Feingold et al., 2019). As there are no previously reported specific markers of bar-shaped SCs, we characterized *I_h_ channel* (*ih*) (Fig. S2A). In addition, we validated *CG30377* for main segment SCs and *α2-adrenergic-like octopamine receptor* (*Octα2R*) for all SCs (Fig. S2B and C). We identified six PC clusters (initial and transitional PCs, main segment PCs, lower tubule PCs, upper ureter PCs, lower ureter PCs, and lower segment PCs). Initial and transitional PCs express *bifid* (*bi*) and *Death executioner Bcl-2* (*Debcl*) (Fig. 1D). The main segment PCs express *urate oxidase* (*Uro*) (Terhzaz et al., 2010, Fig. 1D). The lower segment PC cluster contains three sub-clusters: lower tubule, upper ureter and lower ureter PCs. *Alkaline phosphatase 4* (*Alp4*) is a known marker of lower segment PCs (Yang et al., 2000, Fig. 1D and 1E), upper ureter PCs express *Sex peptide receptor* (*SPR*) (Fig. 1D and 1E), and lower ureter PCs express *Wnt oncogene analog 4* (*Wnt4*) (Fig. 1D and 1E). Thus, marker genes for lower ureter PC is *Wnt4*; upper ureter is *SPR*; lower tubule is *SPR*^-^, *Alp4*^hi^; and the main segment PC is *Uro* (Fig. 1F). We also identified a small cell cluster expressing *Extended synaptotagmin-like protein 2* (*Esyt2*) that we named lower segment PC. Note that we find that *Alp4* is expressed in three clusters (the upper ureter PC, lower tubule PC, and lower segment PC clusters) (Fig. 1D and 1E). *Wnt4*, *SPR*, *Esyt2* and *Debcl* are new marker genes that had not been previously reported.

Nephrocyte clusters are defined by the expression of *sticks and stones* (*sns*), *Cubn* and *prospero* (*pros*). *sns* encodes a core component of slit diaphragm (Zhuang et al., 2009) and *Cubn* encodes a receptor for protein reabsorption (Zhang et al., 2013); both are critical for NC function. These two genes have lower expression in GCs compared to PNs. Two NC-specific marker genes, *Kruppel-like factor 15* (*klf15*) and *UDP-glycosyltransferase family 36 member A1* (*Dot*) (Ivy et al., 2015; Zhang et al., 2013), were not present in our data set, most likely due to technical limitations with the 10X approach as these genes have very short 3’UTRs. *Pros* and *Hand* are known makers for GCs and PNs (Weavers et al., 2009, Fig. S2D and 2E). Details on the marker genes are listed in Supplementary Table 2. Finally, in order to make the dataset accessible to users, we developed a visualization web portal (https://www.flyrnai.org/scRNA/kidney/) that allows users to query the expression of any genes of interest in different cell types.

### Reconciling physiology with clusters

The Malpighian tubule generates a primary urine not by paracellular filtration but by potent active cation transport. This is coupled to channel mediated anion flux to balance charge and water channels to allow rapid flux of osmotically obliged water (Cohen et al., 2020). The tubules can transport their own volume of water every six seconds, making them the fastest-secreting epithelium known (Dow et al., 1994). By contrast with the vertebrate nephron, the paracellular route in Malpighian tubules is tightly guarded by septate junctions, and solutes are excreted by a wide range of massively expressed transporters (Wang et al., 2004; Chintapalli et al., 2007). Many of the genes underlying these processes have been identified and some localized to cell types. However, the single cell dataset allows us to address various questions at a larger scale. In particular, do genes ascribed to particular processes co-locate to the same cell types or regions, what new insights can be gained as to regional specialization, and can we predict functions of previously uncharacterized genes based on their expression patterns?

The PC transcriptome broadly follows expectation (Fig. 2); the plasma membrane V-ATPase subunits all show elevated expression in the PCs along the whole length of the tubule, not just in the main segment (Fig. 2 and Fig. S3A). A candidate apical exchanger, *Na^+^/H^+^ hydrogen exchanger 3* (*Nhe3*), shows a similar expression pattern, as does the Na^+^, K^+^ ATPase that stabilizes cellular cation levels (Torrie et al., 2004) (Fig. 2). This implies that the basic transport machinery is an inherent property of the whole length of the tubule, not just the secretory region. By contrast, the inward rectifier K^+^ channel family genes, all of which are strongly expressed in the tubule, show distinct patterns. *Inwardly rectifying potassium channel 1* (*Irk1*) marks PCs of only the secretory main segment of the tubule (Fig. 2), *irk2* is expressed in the main segment and lower tubule, and *irk3* is generally expressed in PCs (Fig. S3B). Control of transport is clearly critical, and receptors for the three major neuropeptides believed to act on PCs to stimulate secretion are all found in PCs of the main segment; however, their expression patterns are slightly different (Fig. 2). *Capa receptor* (*CapaR*) is confined to the initial, transitional and main segments, as is its effector, the cyclic GMP kinase *foraging* (*for*). *DH31-R* is similarly expressed but *DH44-R2* is present in the main segment and lower tubule. Surprisingly, it is also strongly expressed in SCs; this dual control of two cell types had not been predicted experimentally.

**Figure 2.**
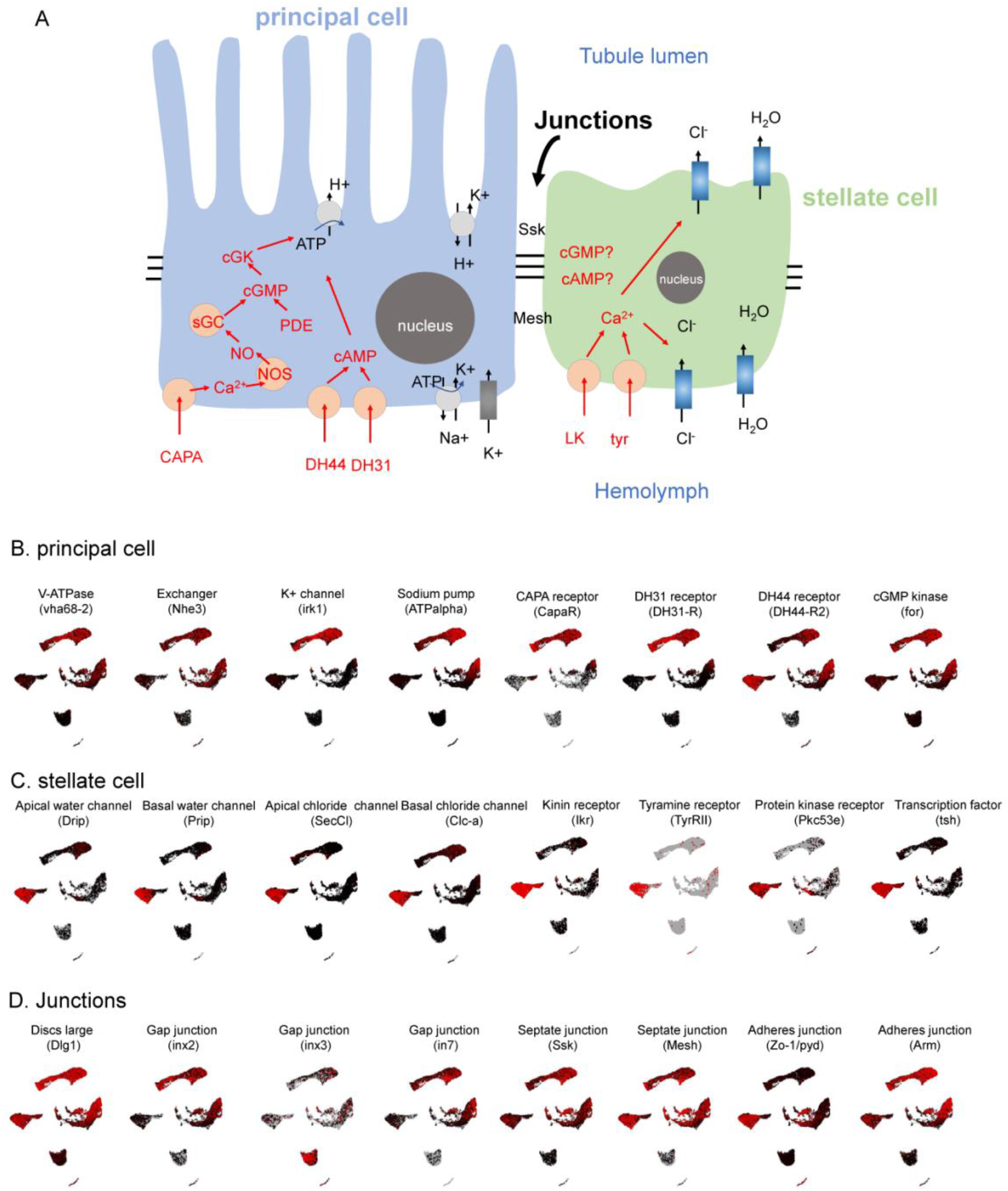
Mapping function to cell types and regions in the tubule. (A) Overview of the two-cell model of insect tubule fluid secretion and its control. Adapted from Dow et al. (2021). (B-D) UMAP distribution of genes involved in principal cell, stellate cell and junctions.

SCs are thought to provide a transcellular shunt for anions and water, and accordingly, the two chloride channels *Clc-a* (Cabrero et al., 2014) and *SecCl* (Feingold et al., 2019), as well as the two true aquaporins *Drip* and *Prip* (Cabrero et al., 2020), show strong SC-enriched expression. Tyramine signals identically to the kinin neuropeptide (Cabrero et al., 2013) and both their receptors show strong localization to SCs, together with their downstream effector, protein kinase C, and the master transcription factor, *teashirt* (*tsh*), which specifies SC fate (Denholm et al., 2013) (Fig. 2). However, there are surprises in this dataset; none of these genes show strong expression in bar-shaped cells, characteristic of the initial segment of anterior tubules, suggesting that although they are developmentally linked to stellate cells, bar-shaped cells are not able to either receive diuretic signals or respond to them.

Junctional permeability is critical in epithelia. As the PC and SC lineages have distinct origins (ectodermal and mesodermal, respectively) (Denholm et al., 2003), they might not necessarily form heterotypic junctions. In fact, both cell types express the septate (occluding) junctional genes *discs large 1* (*dlg1*), *snakeskin* (*ssk*) (Dornan et al., 2020) and *Mesh* (Jonusaite et al., 2020) throughout the length of the tubule, suggesting that both PC-PC and PC-SC junctions are equally tight. Interestingly, although both cell types also contribute adherens junction components, the emphasis is different, with *polychaetoid* (*ZO-1/pyd*) emphasized by SCs and *Armadillo* (*arm*) by PCs. Of the gap junction (innexin) genes, three are strongly expressed in tubule (Chintapalli et al., 2013); *inx2* and *inx7* are expressed in PCs but not SCs, and *inx3* is enriched in stem cells. Thus, PCs have the abilities to communicate (for example, sharing second messengers) and synchronize activities, but SCs are likely to be functionally independent. There is experimental evidence to support this idea; stimulation of PCs with *Capa* elevates intracellular calcium in PCs but not SCs (Rosay et al., 1997), whereas the opposite holds for *Kinin* signaling (Radford et al., 2002). The *Capa* and *Kinin* pathways thus act independently on two cell types without detectable crosstalk, making functional interaction unlikely (MacMillan et al., 2018).

The tubules show strongly enriched expression of most organic solute transporter families (Wang et al., 2004), including the ABC-transporters that underly eye color (*white* (*w*), *scarlet* (*st*), *brown* (*bw*)), and these are all confined to main segment PCs (Fig. S3E). Tubules are also excellent models for renal diseases (Cohen et al., 2020; Dow and Romero, 2010) and readily develop oxalate kidney stones. Knockdown of the oxalate transporter *prestin* increases these stones, presumably by preventing reuptake of secreted oxalate (Hirata et al., 2012; Landry et al., 2016); *prestin* is expressed in PCs of the reabsorptive (O’Donnell and Maddrell, 1995) lower tubule (Fig. S3E). Similarly, transporters that have been implicated in the excretion of xenobiotics (Torrie et al., 2004) are expressed only in PCs (Fig. S3F), confirming the role of these cells in general-purpose solute transport.

As well as transport, tubules play a liver-like role in detoxification, and show conspicuous expression of genes shown to detoxify insecticides (e.g. *Cyp6g1* and *Cyp12d1*, Catania et al., 2004; Yang et al., 2007; Le Goff et al., 2003), and the master transcriptional regulator *Hr96* (King-Jones et al., 2006); all of these genes show close co-expression in PCs (Fig. S3G). Several transcription factors allow the clusters imputed here to be resolved. For example, *tsh* and *tiptop* (*tio*) are SC-specific, *N* marks stem cells, and *dachshund* (*dac*), *Dorsocross1* (*Doc1*), Homothorax (*Hth*), and *cut* (*ct*) provide graded resolution of PC domains (Fig. S3K-M).

### Control of stellate cell shape

SCs, which control channel-mediated Cl^-^ and water flux, have a cuboidal shape in third instar larvae. Subsequently, during the pharate adult stage, they adopt a star shape in the main segment and a bar shape in the initial segment (Beyenbach et al. 2010; Cabrero et al., 2020; Dow, 2012; Choubey et al., 2020). Previous studies have shown that disruption of SCs affects fly survival. For example, conditional downregulation of *Rab11* in SCs results in lethality at the pharate adult stage, and knockdown of *Snakeskin* (*Ssk*) in SCs result in loss of fluid integrity and a significant reduction in viability (Choubey et al., 2020; Dornan et al., 2020). Furthermore, ablation of SCs causes lethality, confirming the essential role of this cell type (Denholm et al., 2003). We identified two sub-clusters of SCs, bar-shaped SCs and main segment SCs (Fig. 3A). GO analysis revealed that bar-shaped SCs play important roles in cell-cell adhesion and potassium transport, while main segment SCs are mainly involved in water, hormone and neuropeptide flux (Fig. S4, Supplementary Table 4).

**Figure 3.**
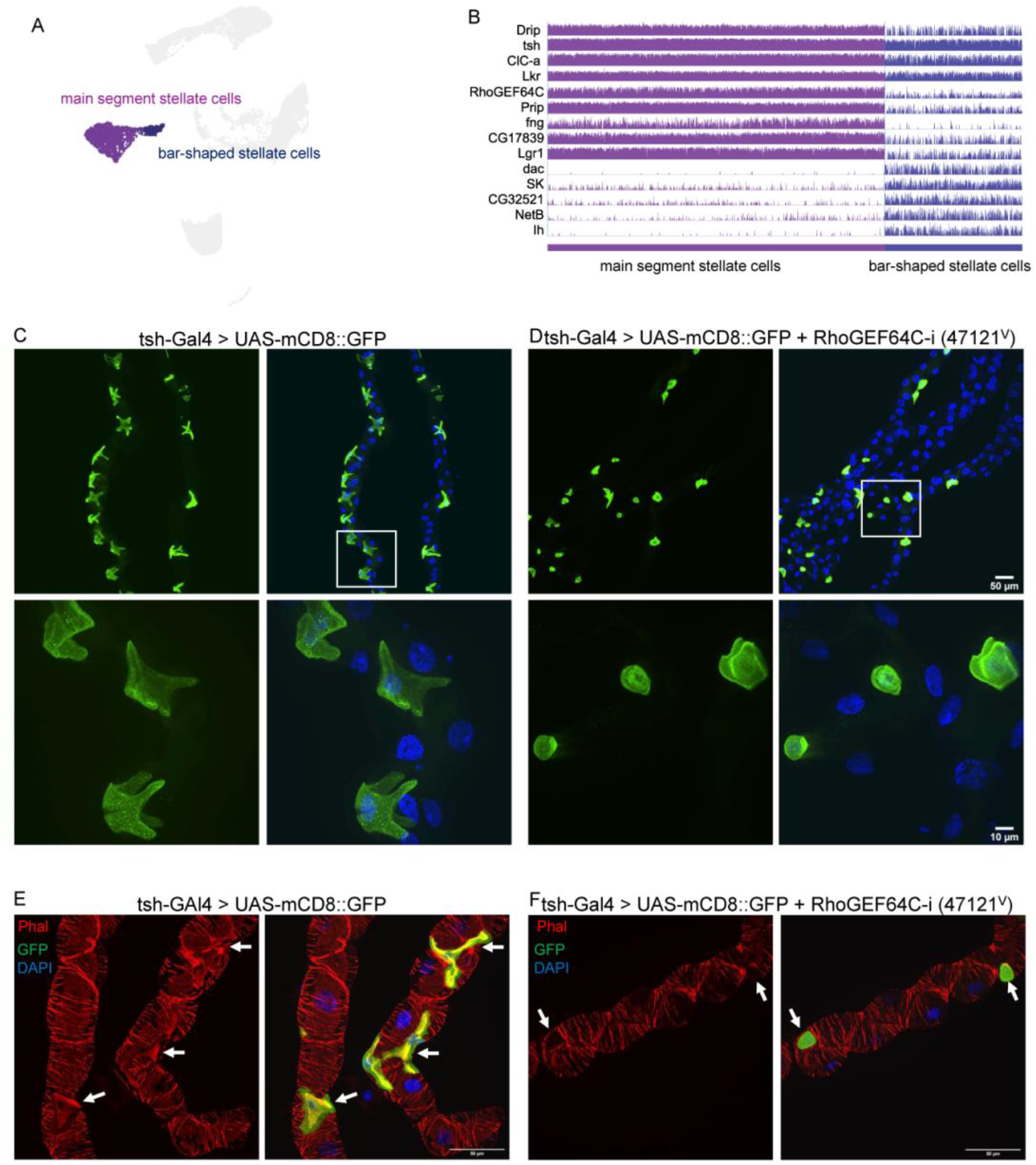
*RhoGEF64c* maintains stellate cell shape. (A) UMAP of two sub-clusters of SCs in the Malpighian tubules. (B) Gene expression level in the population of all SCs. (C) Cell shape visualized using tsh-Gal4 driving mCD8-GFP. DAPI (blue) is used to stain nuclei. White box indicates the zoom-in region. (D) Knockdown using VDRC line 47121^v^ of *RhoGEF64c* affects SC cell shapes. (E and F) Knockdown of *RhoGEF64c* results in loss of cytoarchitectural organization. Cell cytoarchitecture is visualized by Phalloidin (Phal; F-actin) staining. Arrows indicate SCs. Scale bars = 50 μm.

Next, we knocked down the top SCs marker genes using *tsh-Gal4* to study their functions, focusing on SCs shape and viability (Fig. S5A). Among the 18 genes analyzed, 13 were associated with reduced viability, four affected main segment SC cell shape, and two reduced main segment SC cell number (Fig. S5B-F). Among these genes, the top-ranking marker gene, *Rho guanine nucleotide exchange factor at 64C* (*RhoGEF64C*) (Fig. 3B), is an exchange factor for Rho GTPases. Knocking down *RhoGEF64c* affects cell shape of the main segment SCs (Fig. 3C and 3D) and viability (Fig. S7B). In humans, Rho-GTPases regulate the formation and maintenance of long cellular extensions/foot processes and their dysfunctions are associated with nephrotic syndrome (NS) (Matsuda et al., 2021). Further, following podocyte injury, Rho-GTPases orchestrate the rearrangement of the actin cytoskeleton (Matsuda et al., 2021). Interestingly, knockdown of *RhoGEF64c* results in loss of cytoarchitectural organization in main segment SCs (Fig. 3E and 3F) but did not affect septate junctions (Fig. S6A and S6B). This contrasts with knockdown of *Ssk,* which caused loss of both cytoarchitectural organization and septate junctions (Dornan et al., 2020). Finally, knockdown of other top marker genes, namely *Prip*, *Frequenin 2* (*Frq2*) and *CG10939,* did not affect the septate junctions (Fig. S6C and S6D).

### Developmental trajectory analysis of principal cells

PCs are mitochondria-rich and transport protons through an apical, plasma membrane vacuolar H^+^-ATPase (V-ATPase) (Davies et al., 1996). The main functions of PCs are to set up a potassium gradient (Day et al., 2008; O’Donnell and Maddrell, 1995), which enters the cell basolaterally through the combined activity of Na^+^, K^+^-ATPase (Torrie et al., 2004), inward rectifier potassium channels (Evans et al., 2005; Wu et al., 2015; Swale et al., 2016), and potassium cotransports (Sciortino et al., 2001; Linton and O’Donnell 1999; Rodan et al., 2012). We identified six PC clusters from the scRNA-seq dataset and identified Gal4 lines that allowed us to precisely map their anatomical locations (Fig. 1D). To understand the functional differences of each PC cell cluster, we performed a GO analysis based on marker genes (Fig. S7A). Most of the GO terms refer to transport and responses to toxic substances, reflecting the main functions of the tubule. Interestingly, the top 10 terms in lower tubule PCs refer to transport, suggesting that the function of lower tubule PC is to transport substances between Malpighian tubules and the hemolymph (GO information is in Supplementary Table 5). scRNA-seq enables the exploration of the continuous differentiation trajectory of a developmental process. Thus, to analyze the developmental trajectory of PCs, we conducted a pseudotime analysis by ordering cells along a reconstructed trajectory using Monocle3 (Fig. S7B and S7C). Consistent with the distribution distance on the UMAP, inferred trajectories demonstrated gradual transitions from cells in lower ureter PCs, upper ureter PCs, lower tubule PCs, and lower segment PCs to main segment PCs, initial and transitional PCs (Fig. S7D). On the UMAP, lower segment PCs are close to lower ureter PCs, upper ureter PCs, and lower tubule PCs. The pseudotime analysis also showed that the state of lower segment PCs is a co-mixture of lower ureter PCs, upper ureter PCs, and lower tubule PCs (Fig. S7D). These results are consistent with our observation *in vivo* using *Esyt2-Gal4* flies (Fig. 1D and 1E), which suggested that lower segment PCs represent a new cell cluster that is different from previously reported *Alp4* expressed cells.

We chose *Best2*, *bifid (bi), Sarcoplasmic calcium-binding protein 2* (*Scp2*), *PDGF-and VEGF-receptor related* (*Pvr*), *Uro*, *salty dog* (*salt)*, *Alp4*, *SPR*, and *Transient receptor potential cation channel A1* (*TrpA1*) as representative genes for each cluster (Fig. S7E). A survey of our scRNA-seq dataset revealed that the expression of *TrpA1* gradually decreased along the pseudotime, followed by increased transcription of *Alp4* and *SPR*. The expression of *Pvr*, *Uro* and *salt* was elevated at the more geographical distant region of main segment PCs, while the progressive increase of *Best2*, *bi*, and *Scp2* expression was only observed in initial and transitional PCs (Fig. S7E). These results indicate that the patterns of expression of marker genes in each cluster are in concordance with the pseudotime analysis of the clusters.

### Cell-type-specific expression of transcription factors and regulatory landscape

To investigate transcription factors (TFs) that may contribute to kidney differentiation and function, we identified 44 cell-type-specific transcription factors by setting up the parameter cutoff based on gene expression levels (fold change > 3) and adjusting the p-value (<0.05) (Fig. 4A-E). We also applied SCENIC, which is designed to reveal TF-centered gene co-expression networks (Aibar et al., 2017), for the simultaneous reconstruction of gene regulatory networks and identification of cell states (Fig. 4F, detail genes name in Fig. S8). By inferring a gene correlation network followed by motif-based filtration, SCENIC keeps only potential direct targets of each TF as modules (regulons).

**Figure 4.**
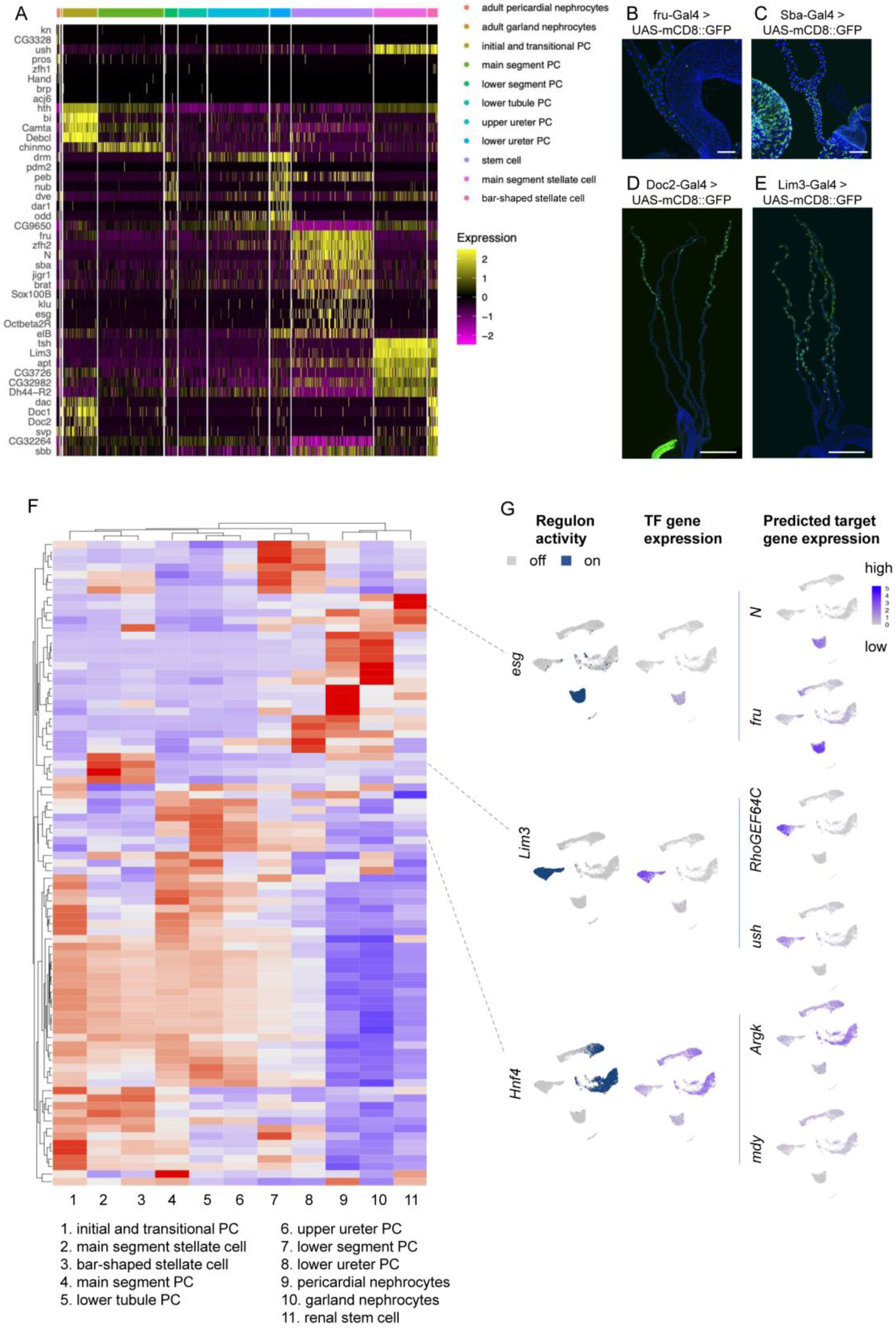
Cell type specific gene regulatory landscape of the fly kidney. (A) Heat map profile of the transcription factors (TFs) in all clusters. Genes were ranked based on expression levels (fold change > 3) and adjusted p-value (<0.05) in each condition in the heat map. (B-E) Expression of *fru*, *Sba*, *Doc2* and *Lim3* visualized using fru-Gal4, Sba-Gal4, Doc2-Gal4 and Lim3-Gal4 driven UAS-CD8::GFP expression, respectively. Scale bars = 100 μm in B and C. Scale bars = 500 μm in D and E. (F) SCENIC results of the fly kidney. The heatmap shows the gene expression level in each cluster. Low regulon activity is shown with blue color and high regulon activity is shown in red. See Sup. FigS8 for an enlarged version of the heat map with gene names. (G) UMAP depiction of regulon activity (“on-blue”, “off-gray”) and TF gene expression (blue scale) of RSCs (*esg*), SCs (*Lim3*), and PCs (*Hnf4*). Examples of target gene expression of the *esg* regulon (*Notch* (*N*) and *fruitless* (*fru*)), *Lim3* regulon (*RhoGEF64C* and *u-shape*d (*ush*)) and *Hnf4* regulon (*Arginine kinase* (*Argk*) and *midway (mdy*)) are shown in blue.

Among the TFs, *esg*, *klu*, and *Sox100B* are specifically expressed in RCSs, which are essential for RSC proliferation and maintenance (Hung et al., 2020). SCENIC could also successfully infer multiple downstream target genes. For example, among the *esg* target genes are *fruitless* (*fru*)*, N*, *Dl*, *klu* (Fig. 4G, Supplementary Table 6). *fru* is expressed male-specifically in the gonad stem cell (GSC) niche and plays important roles in the development and maintenance of GSCs (Zhou et al., 2021). We found that *fru* is also expressed in RCSs of both sexes (Fig. 4B), suggesting that it plays a critical role in RSC proliferation and/or maintenance.

*tsh*, *tio*, and *Lim3* are expressed in both main segment and bar-shaped SCs (Fig. 4A). *tsh* and *tio* are paralogous genes that control SCs shape and the expression of genes required for terminal physiological differentiation (Laugier et al., 2005; Denholm et al., 2013). Interestingly, human *TSHZ* (homolog of *tsh*) genes are causal kidney disease loci, including ureteral smooth muscle differentiation and congenital pelvi-ureteric junction obstruction (Caubit et al., 2008; Jenkins et al., 2010). In addition, *Dac*, *Doc1* and *Doc2*, which have been associated with tissue morphogenesis (Fig. 4D) (Brás-Pereira et al., 2016; Paul et al., 2018; Fan et al., 2021), are not only expressed in bar-shaped SCs but also in initial and transitional PCs. Finally, SCENIC also reveals that *Lim3* is highly enriched in SCs (Fig. 4E and Fig. S8), which is consistent with the role of *RhoGEF64C* in controlling SC morphology (Fig. 3, Fig. 4G), as *RhoGEF64C* is a predicted target of *Lim3* (Fig. 4G). Altogether, our analysis provides a list of possible TFs that control SCs morphology.

The two pairs of tubules are asymmetric both morphologically and transcriptionally. The anterior (right-side) tubules have an extended initial and transitional segment that typically contains calcareous concretions (stones) (Wessing and Eichelberg, 1978), and show selective expression of *Doc1*, *Doc2* and *dac* (Chintapalli et al., 2012). These genes reflect the initial dorsal specification of the anterior tubules, before an embryonic rotation of the gut places them on the right side (Chintapalli et al., 2012). The single-cell data maps expression of these genes specifically to both bar-shaped cells and PCs of just the initial and transitional segments, suggesting a continuing role in maintaining the unique identities of these cell types. By contrast, PCs of the rest of the tubule, and SCs, show no such expression, implying that they are functionally equivalent in both sets of tubules. One interesting TF expressed in several PCs is *Hepatocyte nuclear factor 4* (*Hnf4*). Its human orthologs are *Hnf4γ* and *Hnf4α,* a major regulator of renal proximal tubule development in mouse (Marable et al., 2020). Purine metabolites, including inosine, adenine, xanthine, hypoxanthine, and uric acid, are associated with increased diabetes risk and diabetic nephropathy, and are increased in *Hnf4* mutant flies (Barry and Thummel, 2016). Interestingly, potential direct targets of *Hnf4* include *Arginine kinase* (*Argk*) and *midway* (*mdy*) (Fig. 4E), with *mdy* acting as a repressor of *Hnf4* and HNF4 controlling lipid metabolism in *Drosophila* nephrocytes (Marchesin et al., 2019). Additional information on these TFs can be found in Supplementary Table 7.

### Metabolic pathway analysis

The basic functions of mammalian kidneys include metabolism of carbohydrates, proteins, lipids and other nutrients. As in mammalian kidneys, insect Malpighian tubules and nephrocytes play an essential role in the maintenance of ionic, acid–base and water balance, and elimination of metabolic and foreign toxins and homeostasis. To further understand metabolism in the fly kidney, we analyzed the KEGG metabolic pathways in UMAP of fly kidney snRNA-seq using AUCell software (Aibar et al., 2017). Among 86 KEGG metabolic pathways, purine metabolism, glycerophospholipid metabolism, nicotinate and nicotinamide metabolism, starch and sucrose metabolism were enriched (Fig. S9). Regarding purine metabolism, Xanthine oxidation is a necessary step in the catabolic pathway for purines toward urate, allantoin and urea. Dysfunction of xanthine oxidase/dehydrogenase (XO/XDH) causes build-up of high levels of xanthine and hypoxanthine forming stones in humans and flies (Dent and Philpot, 1954; Ichida et al., 1997; Miller et al., 2013). Human ancestors lost the ability to synthesize a functional urate oxidase due to multiple point mutations in the *Uro* gene, resulting in increased serum and urinary uric acid (UA) levels (Mandal and Mount, 2015). In the UA pathway, humans and flies share some of the same steps. The product of the fly *Uro* gene which catalyzes formation of allantoin from UA (Fig. 5A). Most UA pathway genes are enriched in fly kidney cells, specifically in PCs (Fig. 5B and C). The enzymes that control the last three steps, which are encoded by *rosy* (*ry*), *Uro* and *CG30016,* are highly enriched in main segment PCs, suggesting that the last step occurs in this region (Fig. 5D). *ry* is the homolog of human XDH and loss-function of *ry* is associated with bloating in the lower tubules and formation of stones (Mitchell and Glassman, 1959). Metabolomic analysis of *ry* mutants showed significant changes up to five metabolites away from the metabolic lesion, with large increases in levels of hypoxanthine and xanthine, and undetectable levels of the downstream metabolite UA (Hobani et al. 2012). The product of *CG30016* is predicted to have hydroxyisourate hydrolase activity and to be involved in purine nucleobase metabolism. It will be interesting to see whether this gene also plays a role in maintaining fly urate levels.

**Figure 5.**
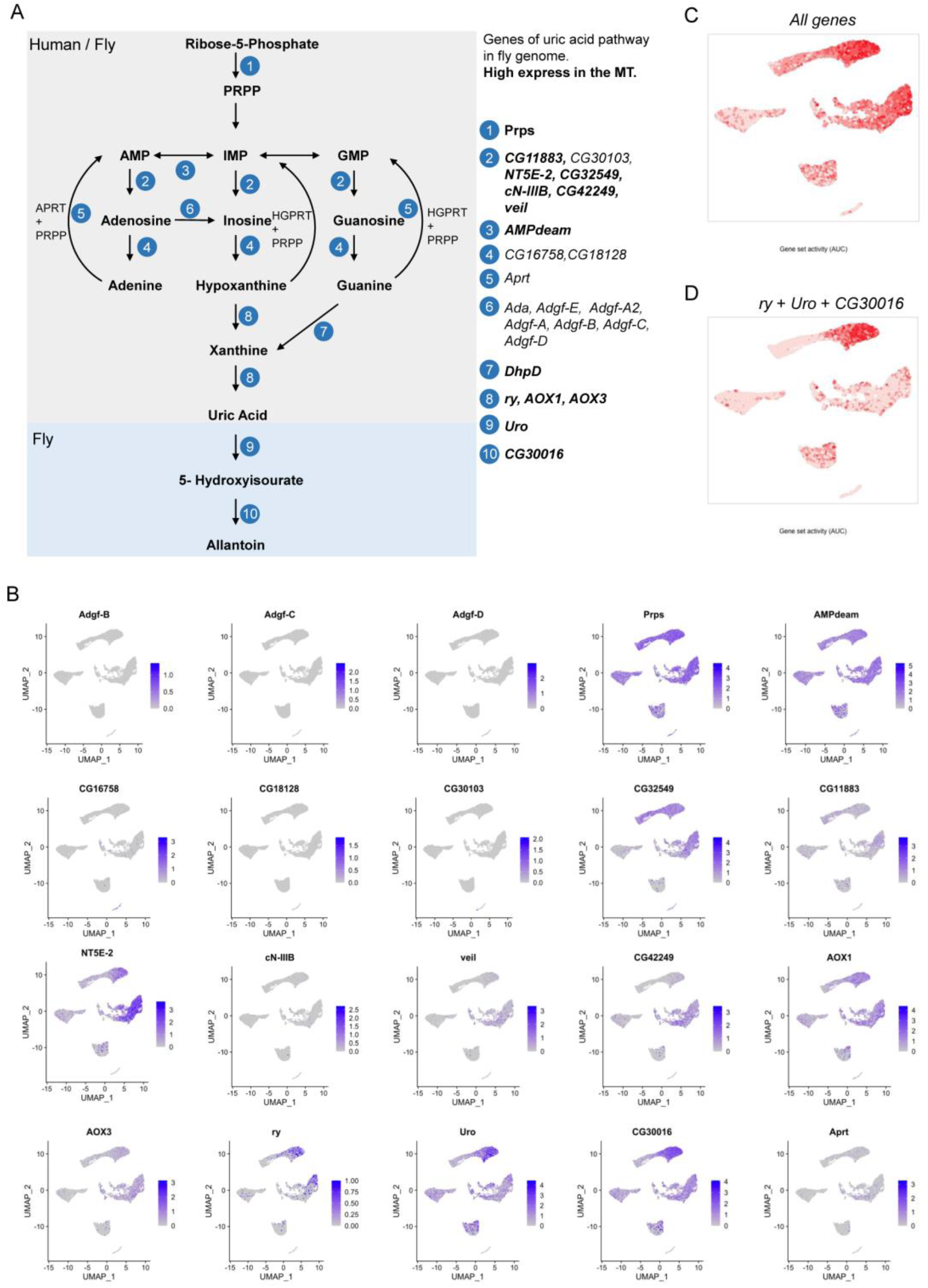
Gene distribution of the Uric Acid pathway in the fly kidney. (A) Uric acid pathway in human and fly. The end product is uric acid in human and allantoin in the fly. Right panel, the enzymes involved at each step. (B) Expression levels of each gene of the uric acid pathway visualized by UMAP. (C) The UMAP plot shows the gene set activity of all genes in the uric acid pathway. (D) Gene set activity of the last three steps, *rosy* (*ry*), *Urate oxidase* (*Uro*) and *CG30016,* visualized by UMAP plots.

Another important function of the insect kidney is detoxification. Many cytochromes P450 (CYPs) genes are involved in this process (Lu et al., 2021) (some are listed in Fig. S10A). For example, *Cyp6g1*, *Cyp6g2*, and *Cyp6A2* are involved in DDT insecticide resistance in flies (Seong et al., 2020; Bergé et al., 1998). *Cyp4e3* has been associated with permethrin insecticide resistance (Terhzaz et al., 2015) and Cyp12a5 in Nitenpyram resistance (Harrop et al. 2018). *Cyp307a2*, *Cyp18a1*, and *Cyp312a1* are involved in degradation of polychlorinated biphenyls (Idda et al., 2020), and *Cyp12d1* impacts caffeine resistance (Najarro et al., 2015). Among these genes, *Cyp6g1*, *Cyp6A2*, *Cyp4e3*, *Cyp12a5*, *Cyp307a2*, and *Cyp12d1* are mainly expressed in PCs (Fig. S10B), which is consistent with PCs playing a key role in detoxification.

### Cross-species and human kidney disease analysis

Considering that the function of all animal excretory systems is to remove toxins from the body and maintain homeostatic balance, we next asked whether we could match fly kidney cell types to higher animals kidney cell types (mouse) and lower animal protonephridia cell types (planarian), and whether the single cell level data can help implicate new genes and cell types in human kidney diseases. We used the Self-Assembling Manifold mapping (SAMap) algorithm (Tarashansky et al., 2021) to map our fly single-cell transcriptomes with scRNA-seq data from mouse (Ransick et al., 2019) and planaria (Fincher et al., 2018). This method depends on two modules. First, a gene-gene bipartite graph with cross-species edges connects homologous gene pairs weighted by protein sequence similarity (all gene pairs are listed in Supplementary Table 8 and Supplementary Table 9). Second, a gene-gene graph projects two single-cell transcriptomic datasets into a joint, lower-dimensional manifold representation, from which each cell mutual cross-species neighbors are linked to stitch the cell atlases together for fly and mouse kidney (Fig. S11A and B). With this method, SAMap produced a combined manifold with a high degree of cross-species alignment (Fig. S11C). After measuring the mapping strength between cell types by calculating an alignment score (as edge width showed in Fig. 6A), which was defined as the average number of mutual nearest cross-species neighbors of each cell relative to the maximum possible number of neighbors, we generated a Sankey plot with 10 fly kidney cell clusters matched to 26 mouse kidney cell clusters (Fig. 6A). A similar analysis was performed for flies and planarians, with 9 fly kidney cell clusters matched to 6 planarian protonephridia cell clusters (Fig. S12A-C and Fig. 6B).

**Figure 6.**
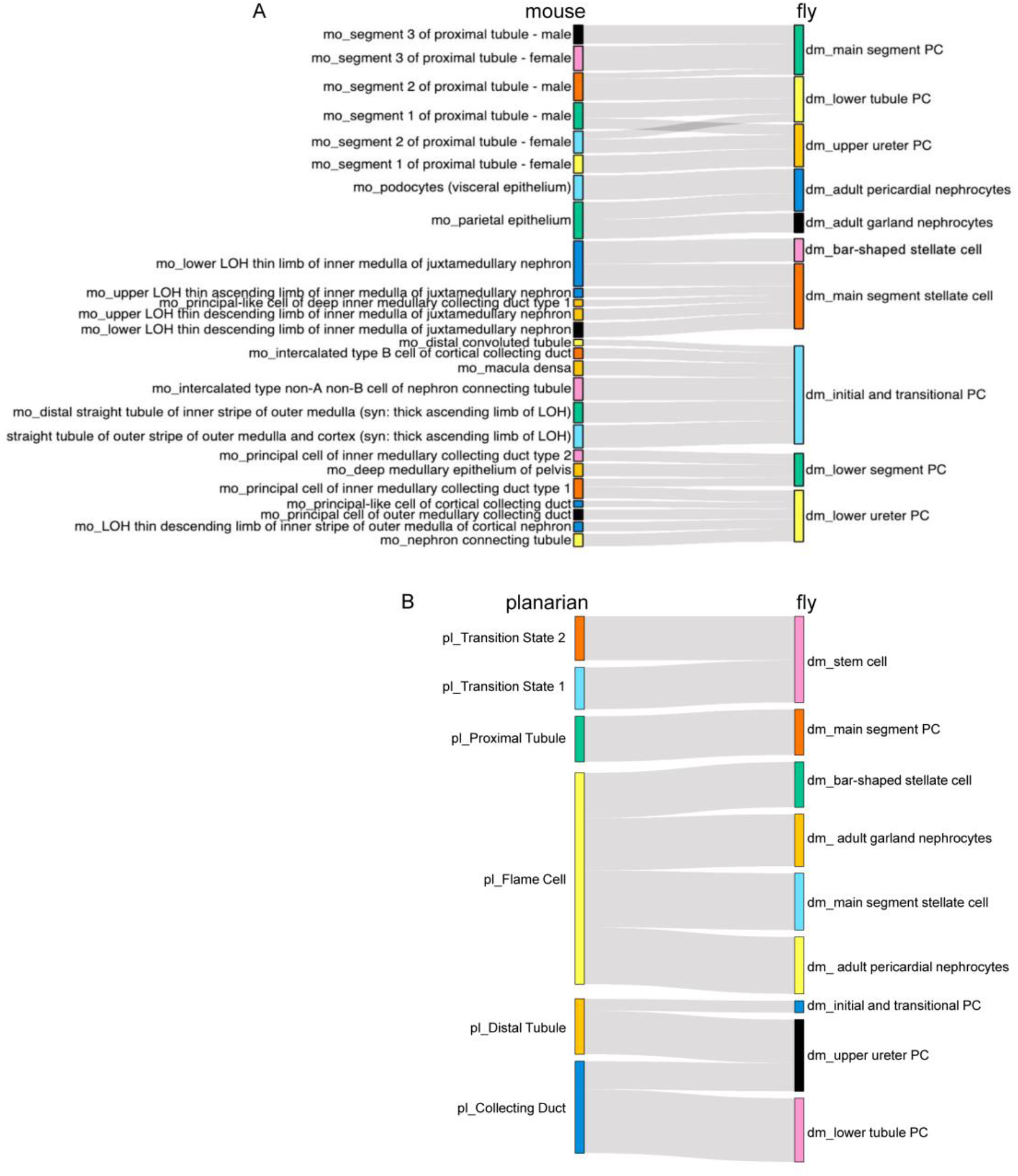
Cross-species analysis of fly, planarian and mouse kidneys using SAMap. (A and B) Sankey plot summarizing the cell type mappings. Edges with alignment scores < 0.1 were omitted.

The results of the fly/mouse analysis suggest that fly main segment PC, lower tubule PC, upper ureter PC are similar to mouse proximal tubule (segment 1-3); fly bar-shaped and main segment SCs map to mouse lower LOH (loop of Henle) thin limb of inner medulla of juxtamedullary nephron; fly adult pericardial nephrocytes are similar to mouse podocytes (visceral epithelium) and parietal epithelium; and fly adult GCs map to mouse parietal epithelium (Fig. 6A). Thus, although pericardial and GC nephrocytes are frequently considered to be interchangeable, they represent different facets of the mammalian nephron. Interestingly, we found that the fly lower segment PCs represents a discontinuous cell population located in the lower segment region and match to mouse PCs of inner medullary collecting duct type 1/2, suggesting that the fly lower segment PC cluster is a newly identified MT cell type.

The fly/planaria comparison suggests that fly stem cells are similar to planarian Transition State 1 and Transition State 2 cluster cells, indicating that kidney stem cells are present in both lower animal species but not mammals. Fly main segment PCs map to planarian Proximal Tubule; fly lower tubule PCs map to planarian Collecting Duct; and fly upper ureter PCs and initial and transitional PCs map to planarian Distal Tubule. Interestingly, fly pericardial nephrocytes, GCs, bar-shaped and main segment SCs map to planarian Flame Cells, suggesting that these cell types have conserved function for removing waste materials (Fig. 6B).

Next, we chose several genes from homologous gene pairs (see Supplementary Table 8 and Supplementary Table 9) to test the robustness of the comparative analyses. Based on the SAMap, fly *pvf1* and *tsh* are highly expressed in SCs. Strikingly, the homologous mouse genes *pdgfa* and *Tshz2* are highly expressed in lower LOH thin limb of inner medulla of juxtamedullary nephron (Fig. S11D). Further, fly *Cyp6g1* and *Na^+^-dependent inorganic phosphate cotransporter* (*NaPi-T*) genes are highly expressed in main segment PC, and their corresponding genes in the mouse, *Cyp4b1* and *Slc22a6*, are highly expressed in mouse proximal tubules (Fig. S11D). *Esyt2* is a marker gene for fly the lower segment PCs, and its homologous gene *Esyt1* is highly expressed in PC of the inner medullary collecting duct type 1/2 (Fig. S11D). In the fly, *sns* is highly expressed in nephrocytes, and the homologous gene in the mouse, *Nphs1,* is highly expressed in mouse podocytes (Fig. S11D). With regards to planaria, despite the lower extent of genome annotation, we identified some informative gene pairs (see Supplementary Table 9) that include the fly nephrocyte marker gene *sns*, the SC marker gene *Nep2*, the stem cell marker gene *esg*, and the main segment PC marker gene *salt*, which could be mapped to the planaria cell clusters (Fig. S12D). Altogether, these results indicate that the SAMap mapping results are well supported by conserved gene expression programs.

Finally, we examined whether the single cell data can help implicate cell clusters and gene targets in human kidney diseases, especially as a previous study in the mouse has shown that hereditary human kidney diseases characterized by the same phenotypic manifestations originate from the same cell types (Park et al., 2018). Strikingly, single cell distribution of human kidney diseases in the fly kidney showed that most of these genes were enriched in the orthologous cell types (Fig. S13). In particular, the fly homologs of 13 of 33 genes associated with monogenic inheritance of nephrotic syndrome in humans were expressed in fly nephrocytes. In the mouse, homologs of genes associated with the syndrome were expressed in podocytes (Park et al., 2018). Among the fly homologs, *sns*, *kin of irre* (*kirre*), and *cubn* have been shown to have key functions in fly nephrocytes (Helmstädter et al., 2017). The fly orthologs of human *Nphs1* and *Kirrel1, sns* and *kirre*, direct adhesion, fusion and formation of a slit diaphragm (SD) structure in insect nephrocytes (Zhuang et al., 2009). Knockdown of *sns* or *kirre* leads to a dramatic decrease in uptake of large proteins, consistent with the role of the SD in mammalian podocytes (Zhuang et al., 2009). Finally, the fly homologs of two genes associated with nephrolithiasis, *ATP6V1B1* and *ATP6V0A4* (*Vha55* and *Vha100-2* in flies), are highly expressed in Malpighian tubule PCs. Mutations in *ATP6V1B1* and *ATP6V0A4* have been identified in calcium oxalate kidney stone patients, suggesting that they are essential for calcium oxalate kidney stone formation (Dhayat et al., 2016). In the mouse, the orthologs of these two genes are hallmarks of intercalated cells, and one type of intercalated cell (intercalated type non-A non-B cell of nephron connecting tubule) matched with fly initial and transitional PCs (Fig. 6A). Interestingly, flies with knockdown of *Vha55* or *Vha100-2* in the Malpighian tubule also develop calcium oxalate kidney stones (Fan et al., 2020). Altogether, the analysis of the expression of fly homologs of human kidney disease-associated genes at the single cell level will help develop more accurate fly models of human kidney diseases.

## DISCUSSION

Here, we surveyed the cell types of the adult fly kidney using snRNA-seq and identified all known cell types and their sub-types. Our dataset provides insights in SC shape, identifying RhoGEF64c as a key cell shape regulator. Interestingly, six clusters of PCs mapped to different regions of the tubule and we could associate them with different physiological functions. The dataset also provides information about potential gene expression networks of transcription factors. Of particular interest, we find that RSCs contain two clusters distinguishable by expression of *Dl^+^ klu^−^* and *Dl^−^ klu^+^* (detailed information in Supplementary text, Fig. S14 and Fig. S15), reminiscent of ISC/EB cells in the midgut (Hung et al., 2020). In addition, we used FlyPhoneDB (Liu et al, 2021) to predict ligand–receptor interactions between different cell clusters, a resource that will help analyze communication among kidney cells (detailed information in Supplementary text and Fig. S16). Altogether, this study will facilitate future work on the fly kidney and serve as a resource to understand cell-type identity and physiology.

Our study extends a previous report that performed scRNA-seq of the Malpighian tubules with a focus on the ureter and lower tubule (Wang and Spradling, 2020). The previous study captured 710 cells that did not include many types of PCs due to the dissection and few SCs, as these oddly shaped cells were likely not captured efficiently using scRNA-seq. Our study using single nuclei rather than single cells overcame this difficulty, and altogether, we successfully recovered 12,166 cells representing 1,730 main segment SCs and 336 bar-shaped SCs. Nevertheless, snRNA-seq did not capture well the GCs, which have double nuclei (Marchesin et al., 2019), or the pericardial nephrocytes, which have large nuclei, as they were underrepresented in our dataset - an issue also reported in several studies of the mouse kidney (Wu et al., 2019; Ransick et al., 2019). We identified six sub-clusters of PCs that map to different anatomical locations. Importantly, GO analysis showed that PCs in different geographic locations have functional differences. For example, main segment PCs respond to toxic substances, and consistent with this, genes related to insecticide metabolism and genes encoding the last three steps enzymes of the uric acid pathway were enriched in these cells (Fig. 5D and S10). The function of lower tubule PCs relates to transport of different substances between Malpighian tubules and hemolymph (Fig. S7). Altogether, PCs in different locations have distinct physiological functions, highlighting that coordination of PC function is required for Malpighian tubules to perform their normal function.

The cross-species analysis not only provided information about the potential functions of unknow cell types, but also gave us a better comparative understanding of kidney cells from lower species (planaria) to higher species (mouse). For example, fly lower segment PCs map to mouse PCs of inner medullary collecting duct type 1/2 (Fig. 6A), but there is no corresponding cell type in planaria. Results of the cross-species analysis will facilitate study of the functions of specific cell types found in higher animals using lower species as models.

*Drosophila* Malpighian tubules and nephrocytes have been used successfully to model human kidney diseases. For example, mutations in the vacuolar-type H^+^-ATPase (v-ATPase) subunit genes ATP6V1B1 and ATP6V0A4 in humans have been identified in recurrent calcium oxalate kidney stones (Dhayat et al., 2016) and knockdown of the fly homologs, *Vha55* and *Vha100-2,* using Uro-Gal4 led to increased formation of calcium oxalate stones in Malpighian tubules (Fan et al., 2020). Our fly kidney cell atlas will facilitate disease modeling and analysis. First, it will help narrow down the number of genes to be tested in specific cell types, as our snRNA-seq has identified cell type-specific transcriptomes. Second, as we were able to match cell types between the fly and mouse, we are now able to associate human kidney disease-associated genes with specific fly kidney cell types. This critical information should facilitate the development of more accurate fly models of human kidney diseases.

In conclusion, our dataset provides detailed insights into: fly kidney cell type-specific gene expression patterns, the specific transcription factors in each cell cluster, potential cell-cell communication network, cross-species mapping of fly kidney cell types to mouse and planarian kidney cell types; association of human kidney disease-associated genes with specific cell clusters. We also provide a web-based resource for visualization of gene expression in single cells of the fly kidney.

## METHODS

### Single nucleus isolation and sequencing

*Drosophila* Malpighian tubules and nephrocytes were dissociated to single nuclei as previously described (Li et al. 2021) with a few modifications. Malpighian tubules were dissected under a microscope from 5-7-day old *Drip-Gal4>GFP.nls* male and female adult flies. Nephrocytes were dissected under a fluorescence microscope from 5-7-day old *Dot-Gal4>GFP.nls* male and female adult flies. Ten flies at a time were dissected and samples immediately transferred into 1.5 ml EP tube with Schneider’s medium on ice to avoid exposing the tissues to room temperature for a long period of time. Once 50 flies were dissected, EP tubes were sealed with parafilm and put on dry ice. Next steps involved spraying 100% ethanol to the dry ice near the tube to quickly freeze the sample and storing samples at −80°C for long-term. After dissection, samples were spined down (thaw samples from −80°C) in 100 ul Schneider’s medium using a bench top spinner, medium was discarded, and 100 ul homogenization butter (Li et al. 2021) was added. Subsequently, 900 ul homogenization buffer was added, and 1000ul homogenized sample transferred into the 1 ml dounce. Nuclei were released by 15-20 strokes with loose pestle and 15-20 tight pestle on ice. 1000 ul sample was filtered through 5ml cell strainer (35 um), and then filter sample using 40 um Flowmi into 1.5ml EP tube. Centrifuge for 10 min at 1000 g at 4°C. Resuspend in 1000 ul PBS/0.5% BSA with RNase inhibitor. And filter sample using 40 um Flowmi into a new EP tube. Hoechst 33342 was used to stain nuclei for more than 5 min. Then FACS and collect single nuclei into a tube for 10x Genomics.

Ten thousand nuclei were targeted for each sample when loaded into the Chromium Controller (10X Genomics, PN-120223) on a Chromium Single Cell B Chip (10X Genomics, PN-120262), and processed to generate single cell gel beads in the emulsion (GEM) according to the manufacturer’s protocol (10X Genomics, CG000183). The library was generated using the Chromium Single Cell 3′ Reagent Kits v3.1 (10X Genomics, PN-1000121) and Chromium i7 Multiplex Kit (10X Genomics, PN-120262) according to the manufacturer’s manual. Quality control for constructed library was performed by Agilent Bioanalyzer High Sensitivity DNA kit (Agilent Technologies, 5067-4626) for qualitative analysis. Quantification analysis was performed by Illumina Library Quantification Kit (KAPA Biosystems, KK4824). The library was sequenced on an Illumina NovaSeq system or Nextseq 500 instrument.

### Dataset processing

The quality of the raw sequencing data was checked by FastQC software. The raw sequencing data were processed by cellranger count pipeline to generate the single cell matrix for each sample. The single cell matrix was analyzed by the Seurat package and Harmony was used for batch correction. The Malpighian tubules and nephrocytes samples were processed and clustered separately, and then were merged by filtering out unrelated cell clusters. The processed result of the gene expression matrix was used for the downstream data analysis.

To facilitate mining of the datasets, we developed a visualization web portal (https://www.flyrnai.org/scRNA/kidney/) that allows users to query the expression of any genes of interest in different cell types and to compare the expression of any 2 genes in individual cells. This MT dataset can also be mined at Fly Cell Atlas (https://flycellatlas.org/) along with other datasets generated by the FCA consortium (Li et al. 2021).

### Gene ontology (GO) analysis

Gene ontology (GO) analysis was performed by clusterProfiler. The marker genes identified using Seurat were used for GO analysis. The strength of enrichment was calculated as negative of log10(p-value), which was used to plot the barplot.

### Cross-tissue analysis

Cross-tissue analysis data are from the Malpighian tubules processed dataset and midgut dataset (accession code: GSE120537). Before merging the two datasets, the number of Malpighian tubules cells was downsampled to the same size as the midgut dataset. The top markers of renal stem cells (RSC) were calculated by comparing the RSC with the rest of the merged dataset without intestinal stem cells (ISC). The top markers of ISC were calculated by comparing the ISC with the rest of the merged dataset without RSC. Results of the comparison are visualized on a Venn diagram.

### Pseudotemporal ordering of cells using Monocle3

Processed Malpighian tubules dataset was analyzed using Monocle3 for pseudotemporal ordering. The state representing lower ureter PC was chosen as the starting time point. The Ridge plot was generated by extracting the cell clustering and pseudotime information and then visualized by Seurat RidgePlot function. The gene expression heatmap was generated by merging the cells into bins by the order of pseudotime and visualized by the pheatmap R package.

### Transcription factors enrichment and SCENIC analysis

The top markers for each cluster were used as the candidates for transcription factors enrichment analysis. The markers were filtered by fold change > 3 and adjusted p-value < 0.05. The transcription factors from the filtered markers were visualized by the Seurat DoHeatmap

The analysis of regulon activity was conducted using the SCENIC pipeline (Aibar et al., 2017). Cells from the previously processed dataset were selected as the input cells. The TF and co-expressed genes were constructed by GRNBoost2. The TF co-expression gene sets were filtered by the RcisTarget fly database. The regulon activity score AUC (Area Under the Curve) was calculated by AUCell, and the active regulons were determined by the AUCell default parameters. The regulon activity was visualized by the average AUC score for each cluster.

### Cell-cell communication analysis

The cell-cell communication analysis was performed using FlyPhoneDB (Liu et al., 2021). The previously processed gene expression matrix and cell clustering information were used as the input for the analysis. Ligand-receptor interaction scores and specificity were then calculated. Cell communication at the signaling pathway level was visualized by a circle plot. The interaction of ligand-receptor pairs between two cell types was visualized by dot plot. The network was generated based on the MIST database and TF2TG literature using Cytoscape (Hu et al., 2018; Otasek et al., 2019).

### Cross-species analysis

Datasets included in the cross-species analysis were the processed dataset from this study for the fly, the mouse kidney dataset GSE129798 and the planarian dataset GSE111764. The analysis was conducted using the SAMap software (Tarashansky et al., 2021). The input h5ad file for SAMap was processed by the Self-Assembling-Manifold (SAM) algorithm. Alignments for each cell type in fly and mouse were calculated by the get_mapping_scores function. Enriched gene pairs from the aligned cell types were retrieved by find_all function with a default alignment score threshold of 0.1. The SAMap results were visualized by the sankey_plot function.

### Fly genetics

Fly husbandry and crosses were performed under a 12:12 hour light:dark photoperiod at 25°C. *Hand-GFP; 4xHand-Gal4/CyO* and *Dot-Gal4* stocks are gifts from Dr. Han Zhe. *esg-Gal4* and *Pros-Gal4* are from the Perrimon lab stock collection.

The following strains were obtained from the Bloomington *Drosophila* Stock Center: *Drip-Gal4* (BL66782), *UAS-GFP.nls* (BL4776), *UAS-mCD8::RFP* (BL32219), *UAS-mCD8::GFP* (BL32185), *c42-Gal4* (BL30835), *Uro-Gal4; tubGal80ts* (BL91415), *Alp4-Gal4* (BL30840), *SPR-Gal4* (BL84692), *wnt4-Gal4* (BL67449), *Esyt2-Gal4* (BL77712), *Debcl-Gal4* (BL81163), *ih-Gal4* (BL76162), *CG30377-Gal4* (BL67426), *Octα2R-Gal4* (BL67637), *tsh-Gal4* (BL3040), *fru-Gal4* (BL30027), *Sba-Gal4* (BL67640), *Doc2-Gal4* (BL26436), *Lim3-Gal4* (BL67450), *Pvf1-Gal4* (BL23032), *y v; UAS-LucRNAi, attp2* (BL31603), *y v; UAS-wat-RNAi, attp40* (BL67801), *y v; UAS-tutl-RNAi, attp40* (BL54850), *y v; UAS-axed-RNAi, attp40* (BL62928), *y v; UAS-prip-RNAi, attp40* (BL50695), *y v; UAS-prip-RNAi, attp2* (BL44464), *y v; UAS-nep2-RNAi, attp40* (BL61902), *y v; UAS-CG13323-RNAi, attp40* (BL53969), *y v; UAS-Lgr1-RNAi, attp2* (BL51465), *y v; UAS-Octα2R-RNAi, attp2* (BL50678), *y v; UAS-CG30377-RNAi, attp40* (BL51386), *y v; UAS-notum-RNAi, attp40* (BL55379), *y v; UAS-TkR99D-RNAi, attp2* (BL27513), *y v; UAS-ih-RNAi, attp40* (BL58089), *y v; UAS-ncc69-RNAi, attp2* (BL28682), *y v; UAS-frq2-RNAi, attp2* (BL28711), *y v; UAS-qvr-RNAi, attp40* (BL58061), *y v; UAS-CG10939-RNAi, attp40* (BL65156), *y v; UAS-CG42594-RNAi, attp2* (BL35006), *y v; UAS-RhoGEF64c-RNAi, attp2* (BL31130).

The following strains were obtained from the Vienna *Drosophila* Resource Center: *y w (1118)*; *attp landing site* (v60100), *UAS-RhoGEF64c-RNAi* (v47121), *UAS-RhoGEF64c-RNAi* (v105252).

For the screen shown in Fig. S5B, 8 *tsh-Gal4/CyO; UAS-CD::GFP* virgin females were crossed with 4 RNAi males. Flies were raised at 22°C and the ratio of Cy+/Cy was determined.

### Immunostaining and confocal microscopy

*Drosophila* Malpighian tubules (and guts), PNs (included in the whole abdomen) and GCs (and foreguts and crops) from adult females were fixed in 4% paraformaldehyde in Phosphate-buffered saline (PBS) at room temperature for 1 hour, incubated for 1 hour in Blocking Buffer (5% normal donkey serum, 0.3% Triton X-100, 0.1% bovine serum albumin (BSA) in PBS), and stained with primary antibodies overnight at 4°C in PBST (0.3% Triton X-100, 0.1% BSA in PBS). Primary antibodies and their dilutions used were: mouse anti-GFP (Invitrogen, A11120; 1:300) and mouse anti-discs-large (DSHB, 4F3,1:50). After primary antibody incubation, the tissues were washed 3 times with PBST, stained with 4′,6-diamidino-2-phenylindole (DAPI) (1:2000 dilution), Phalloidin TRITC (Sigma-Aldrich, 1:2000) and Alexa Fluor-conjugated donkey-anti-mouse (Molecular Probes, 1:1000), in PBST at 22°C for 2 hours, washed 3 times with PBST, and mounted in Vectashield medium.

All images presented in this study are confocal images captured with a Nikon Ti2 Spinning Disk confocal microscope. Z-stacks of 15-20 images covering one layer of the epithelium from the apical to the basal side were obtained, adjusted, and assembled using NIH Fiji (ImageJ), and shown as a maximum projection. Details of the imaging method are as follows: Samples were imaged with a Yokogawa CSU-W1 single disk (50 µm pinhole size) spinning disk confocal unit attached to a fully motorized Nikon Ti2 inverted microscope equipped with a Nikon linear-encoded motorized stage with a Mad City Labs 500 µm range Nano-Drive Z piezo insert, an Andor Zyla 4.2 plus (6.5 µm photodiode size) sCMOS camera using a Nikon Plan Apo 60x/1.4 NA DIC oil immersion objective lens with Cargille Type 37 immersion oil (cultured cells) or a Nikon Plan Apo 20x/0.75 DIC air objective lens (tissue samples). The final digital resolution of the image was 0.109 and 0.325 µm/pixel, respectively. Fluorescence from DAPI, Alexa Fluor (AF)-488, and AF555 was collected by illuminating the sample with directly modulated solid-state lasers 405 nm diode 100 mW (at the fiber tip) laser line, 488 nm diode 100 mW laser line, and 561 nm DPSS 100 mW laser line in a Toptica iChrome MLE laser combiner, respectively. A hard-coated Semrock Di01-T405/488/568/647 multi-bandpass dichroic mirror was used for all channels. Signal from each channel was acquired sequentially with hard-coated Chroma ET455/50, Chroma ET525/36 nm, and Chroma ET605/52 nm emission filters in a filter wheel placed within the scan unit, for blue, green, and red channels, respectively. Nikon Elements AR 5.02 acquisition software was used to acquire the data. 2 µm range Z-stacks, set by indicating the middle focal plane and a z-step interval of 50 µm, were acquired using piezo Z-device, with the shutter closed during axial movement. Images were acquired by collecting the entire Z-stack in each color or by acquiring each channel in each focal plane within the Z stack. Data were saved as ND2 files.

### Data availability

The authors declare that all data supporting the findings of this study are available within the article and its supplementary information files or from the corresponding author upon reasonable request. Raw snRNA-seq reads have been deposited in the NCBI Gene Expression Omnibus (GEO) database under accession codes: (XXX, will provide soon). Processed datasets can be mined through a web tool (https://www.flyrnai.org/scRNA/kidney/) that allows users to explore genes and cell types of interest.

## Supporting information

These zip contain Supplementary Table 1-12.

## ACKNOWLEDGMENTS

We thank the assistance provided by the Microscopy Resources on the North Quad (MicRoN) core at Harvard Medical School. We thank Sudhir Gopal Tattikota for suggestions on sequencing and data analysis and Stephanie Mohr for comments on the manuscript. Relevant grants support include NIA R00 AG062746 (H.L.), NIDCD R01 DC005982 (LL), NIDDK (DK107350, DK094526, DK110792) (A.P.M.), and BBSRC-NSF (NP). H.L. is a CPRIT scholar. S.R.Q. is an investigator of Chan Zuckerberg Biohub. J.A.T.D. is supported by UK BBSRC grants BB/P024297/1 and BB/V011154/1. L.L. and N.P. are investigators of Howard Hughes Medical Institute.

## AUTHOR CONTRIBUTIONS

N.P. and J.X. conceptualized and designed the experiments. J.X. performed most experiments. Y.F.L. performed the bioinformatics analysis. H.J.L. and S.S.K. performed single nucleus isolation and RNA library construction. R.J.H. helped with dissection experiments and annotation. Y.H. and A.C. helped with bioinformatics analysis and the website. T.A., C.K. and B. W. contributed to cross-species analysis. S.R.Q. and L.L. contributed reagents and supervision of single nucleus isolation and RNA library construction. J.X., Y.F. L., J. A.T.D., A.P.M. and N.P. analyzed the data. J.X. wrote the first draft of the paper. N.P. and J.A.T.D. edited the paper. All authors discussed the results and commented on the paper.

## Supplemental Information

### Supplementary text

#### Similarities between renal and intestinal stem cells

Compared to a previous report that performed scRNAseq of the Mapighian tubules focusing on the stem cell zone (Wang and Spradling, 2020), our study captured more cells and more cell types. Wang and Spradling focused on the response of RSCs to tissue injury. RSCs were previously identified as a distinct population that expresses *esg* (Singh et al., 2007). RSCs are located in the lower ureter, the upper ureter and lower segment of the MT with small nuclei (Takashima et al., 2013). They respond to tissue injury by upregulating the JNK, EGFR/MAPK, Hippo/Yki and JAK/STAT pathways that promote RSC daughter differentiation (Wang and Spradling, 2020). RSCs originate from the same pool of adult midgut progenitors that generate the posterior midgut intestinal stem cell (ISCs) (Takashima et al., 2013; Xu et al., 2018). To examine how similar RSCs are to ISCs, we compared the snRNA-seq RSC data with previously reported scRNAseq ISC data (Hung et al., 2020). Consistent with their common origin (Takashima et al., 2013), the two stem cell clusters have high similarity at the gene expression level compared to other cell clusters (Fig. S14A).

The *esg* gene, a stem cell marker for both RSCs and ISCs (Fig. S14B and 14C), encodes a transcription factor that contributes to stem cell maintenance through modulation of Notch activity (Loza-Coll et al., 2014). In the intestine, *esg* is not only expressed in ISCs but also in AstC-EEs (enteroendocrine cells that express *Allatostatin C*, *AstC*) and NPF-EEs (EEs that express *neuropeptide F*, *NPF*) (Hung et al., 2020). In the intestine, ISCs are highly mitotic, especially during regeneration, and give rise to a transient progenitor, the enteroblast (EB) (Ohlstein and Spradling, 2006; Micchelli and Perrimon, 2006), whereas in the Malpighian tubule, RSCs normally divide very slowly (Wang and Spradling, 2020). Interestingly, 56 genes overlapped between RSCs and ISCs (MT^+^gut^+^, genes highly expressed in RSCs and ISCs), including *esg*, *N*, *Dl*, *klumpfuss* (*klu*), and *Sox100B* (Fig. S15A, all genes are listed in Supplementary Table 10. Hung et al., 2020). Gene Ontology (GO) analysis reveals that MT^+^gut^+^ genes are enriched in cell differentiation, proliferation, and stem cell division. However, GO terms of MT^+^gut^-^ (genes highly expressed in RSCs, low or not expressed in ISCs) mainly contain genes annotated as involved in growth, tube morphogenesis, and epithelial cell differentiation. Finally, GO terms of MT^-^gut^+^ (genes highly expressed in ISCs, low or no expressed in RSCs) mainly represent genes involved in protein folding, translational initiation, and peptide biosynthesis (Fig. S15B, all GO terms are listed in Supplementary Table 11). Protein folding is relevant to the stress response, reflecting damage to the gut caused by the food. Altogether, these analyses suggest that RSCs and ISCs have a common origin but are in different cell states.

In our previous gut scRNA-seq study, ISCs/EBs were annotated as one cluster based on the expression of *Dl* and *esg*. However, this cluster could be split into ISCs and EBs, as one subset of cells in the ISC/EB cluster is *Dl^+^ klu^−^* and another subset is *Dl^−^ klu^+^* (Hung et al., 2020). Interestingly, we could also identify two RSC sub-clusters based on the expression of *Dl^+^ klu^−^* and *Dl^−^ klu^+^* (Fig. S14D and S14E). The *Dl^+^ klu^−^* sub-cluster specifically expresses *Dl*, *N*, and *esg*, reminiscent to the *Dl^+^ klu^−^* sub-cluster of ISC (Fig. S14F), whereas the *Dl^−^ klu^+^* sub-cluster specifically expresses *E(spl)m3-HLH*, *E(spl)malpha-BFM, E(spl)mbeta-HLH*, which are transcription factors executing Notch-mediated cellular differentiation (Couturier et al., 2019; Lu and Li, 2015, Fig. S14F).

### Cell-cell communication networks in the fly kidney

Previous studies have indicated that the survival, renewal, and differentiation of PCs and SCs are largely regulated through their cross-talk with RSCs (Singh et al., 2007; Takashima et al., 2013; Wang and Spradling, 2020). We used FlyPhoneDB (Liu et al, 2021) to explore cell–cell communication between the different fly kidney cell clusters. FlyPhoneDB was established recently and provides predictions of ligand-receptor interactions based on fly scRNA-seq data. We analyzed 13 major pathways and indicated their cell–cell interaction pairs between the different cell clusters (Fig.S16A). Strikingly, the Notch ligand only has interaction within RSCs and does not pair with other cell clusters (Fig. S16A). This is consistent with previous studies showing that differential Notch activity is required for RSC homeostasis and that damage activates Notch signaling, which in turn regulates differentiation of RSCs to PCs (Li et al., 2014; Wang and Spradling, 2020). Further, we found that the EGFR signaling pathway connects RSCs and all SCs and PCs, with a preferentially strong interaction with main segment SCs and main segment PCs (Fig. S16A). These are consistent with previous studies showing that EGFR is dispensable for RSC maintenance but required for RSC proliferation (Li et al., 2015). In addition, FlyPhoneDB predicts a strong interaction from main segment SCs to main segment PCs with the Pvf1-Pvr ligand-receptor pair (Fig. S16B). This interaction is based on the gene expression pattern in cell clusters of main segment SCs and PCs via MIST database and TF2TG literatures (Fig. S16C). Consistent with this, *pvf1* was highly expressed in main segment SCs as detected using *pvf1-Gal4>mCD8::GFP* (Fig. S16D). Altogether, FlyPhoneDB predicts a number of specific signaling events between Malpighian tubule cell clusters. The full list of predicted gene pairs can be found in Supplementary Table 12.

---

**Figure S1.**
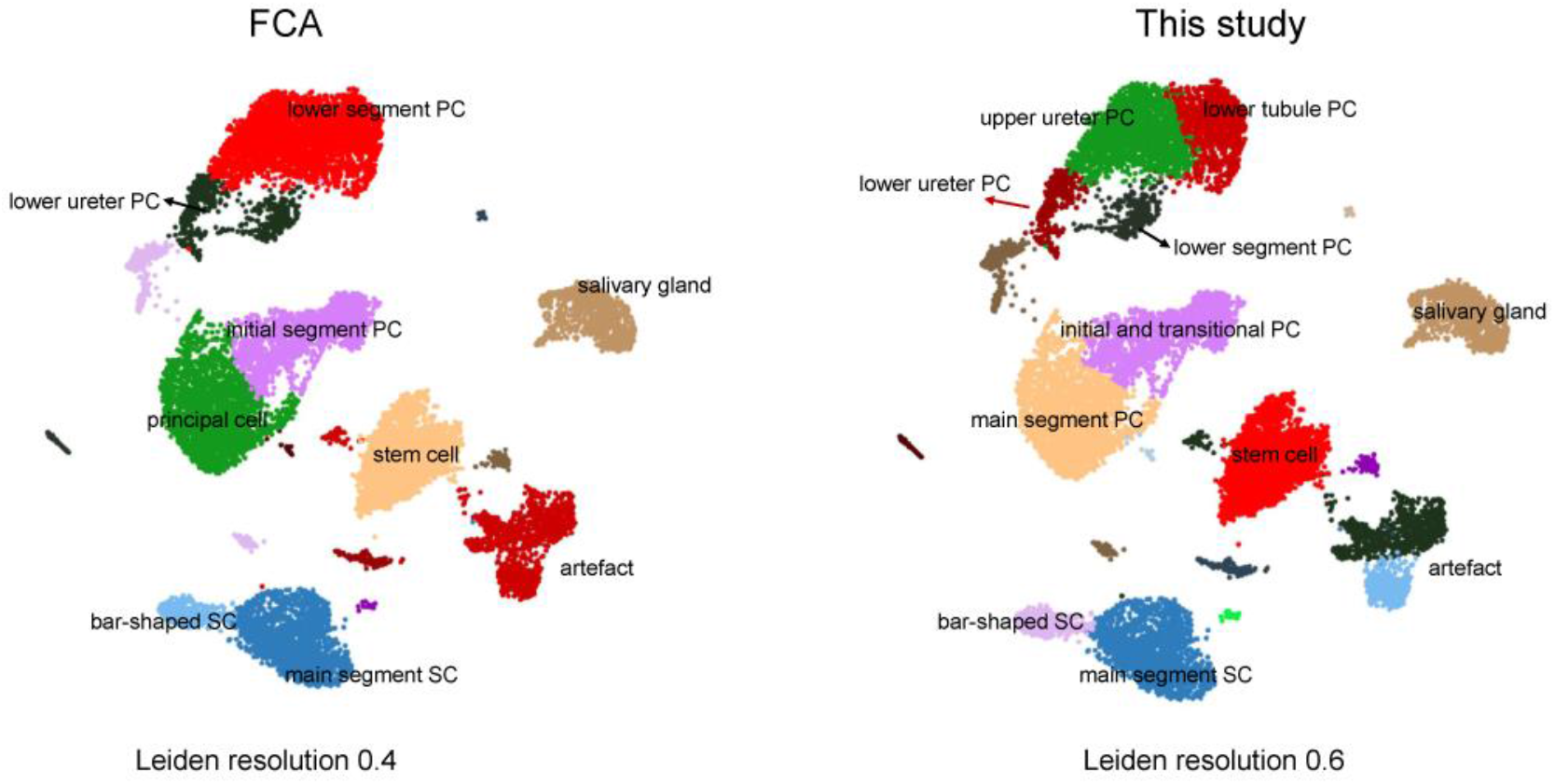
High resolution snRNA-seq analysis of the adult MT. Left, UMAP of the MTs FCA data set at Leiden resolution 0.4 (Li et al., 2021). Right, annotation of the same data set at Leiden resolution 0.6 (this study). The FCA analysis reports four clusters for principal cells: lower ureter PC, lower segment PC, principal cell, and initial segment PC. In this study, we defined six clusters for principal cells based on Gal4 reporter lines: lower ureter PC, lower ureter PC, lower tubule PC, lower segment PC, main segment PC, and initial segment PC. Note that this figure contains all the original clusters, including non-Malpighian tubule cell clusters (salivary glands, artefacts), which we did not include in Fig. 1B.

**Figure S2.**
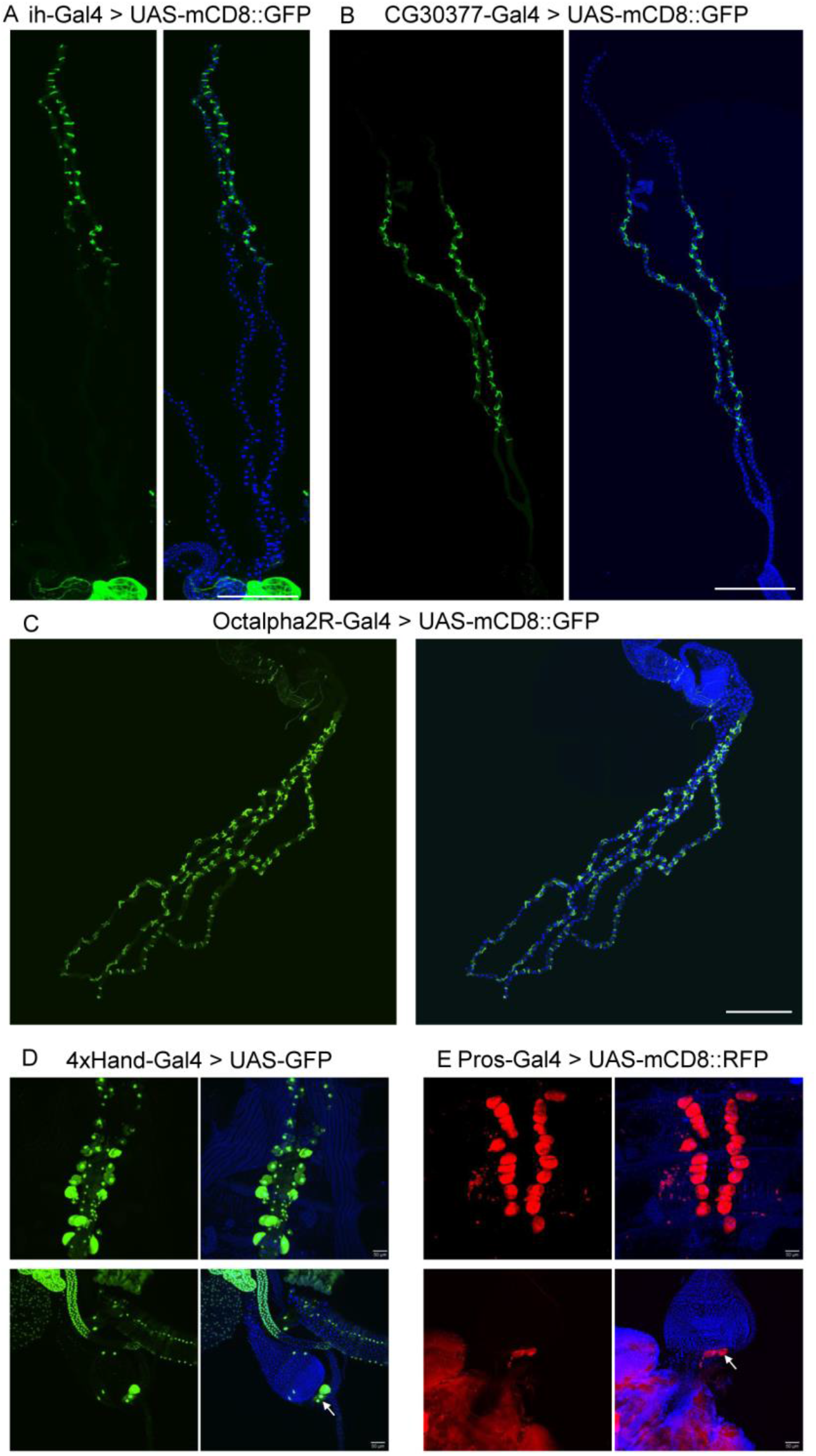
Gene expression in stellate cell clusters and nephrocytes. (A-C) The expression of three new marker genes for SCs is shown using Gal4 lines driving UAS-mCD8::GFP: *I_h_ channel* (*ih*) (bar-shaped SC), *CG30377* (main segment SCs) and *Octalpha2R* (all SCs). Scale bars = 500 μm. (D and E) *Hand* and *Prospero (Pros)* genes are previously known marker genes. *4xHand-Gal4* line is a four copy enhancer sequences of the *Hand* gene driving Gal4 (Zhu et al., 2017). Arrows indicate GCs. Scale bars = 50 μm.

**Figure S3.**
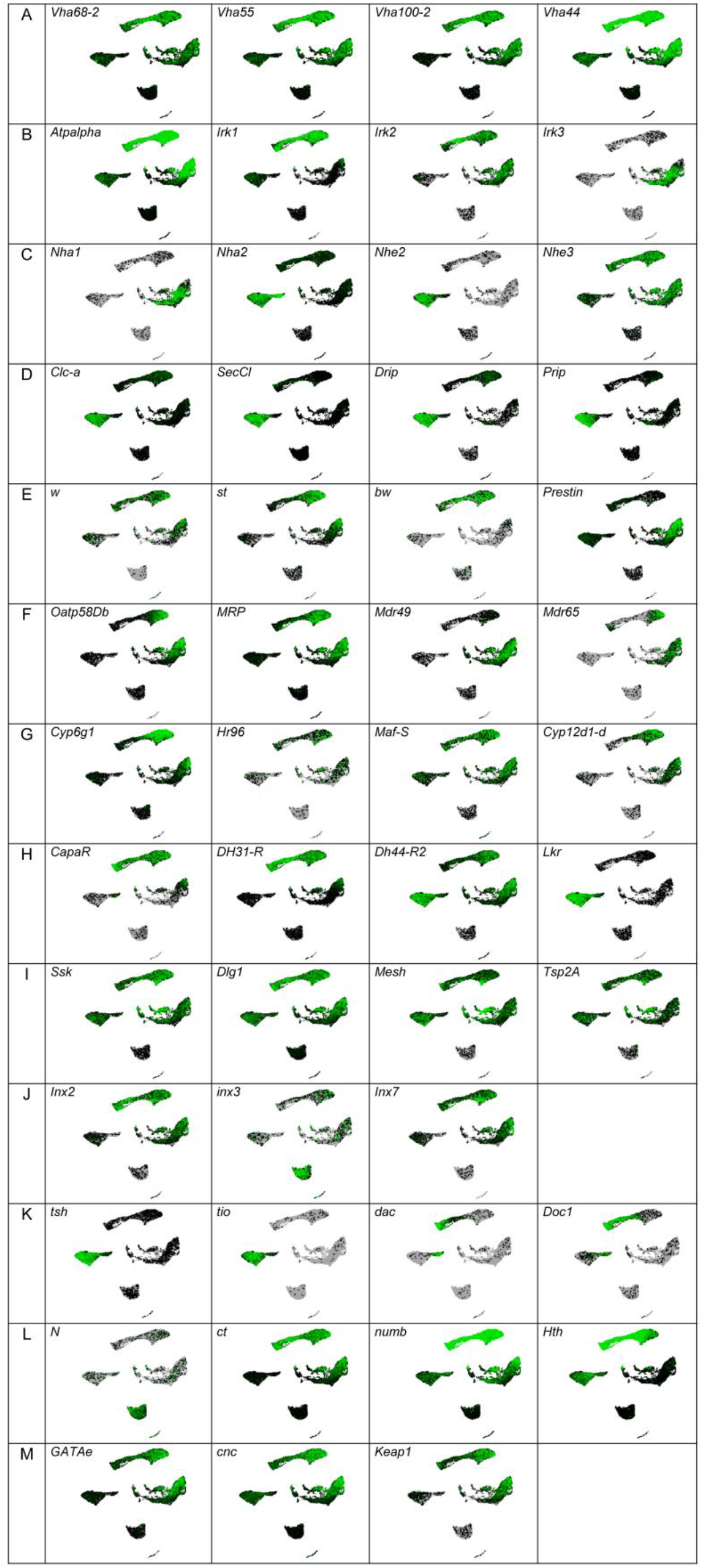
Expression encyclopedia of function-linked tubule genes.

**Figure S4.**
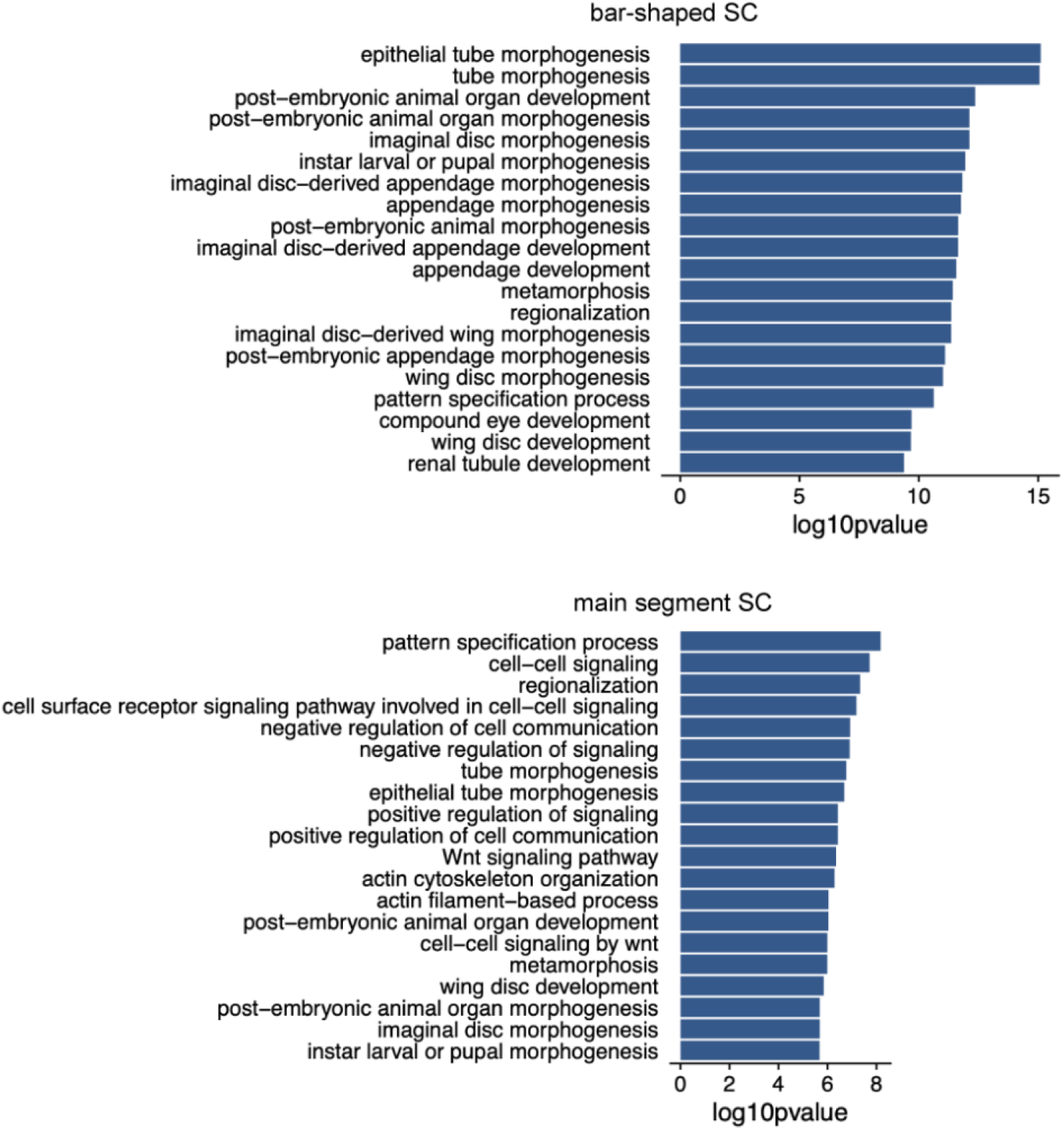
Gene Ontology (GO) analysis of bar-shaped SC and main segment SC clusters. Bar-shaped SCs are enriched for epithelial tube morphogenesis and tube morphogenesis GO terms. These cells specifically express the cell adhesion genes *turtle* (*tutl*), *Tenascin accessory* (*Ten-a*) and *echinoid* (*ed*), the transcription factors *dachshund* (*dac*), *Dorsocross2* (*Doc2*) and *Doc1*, and the potassium channels *tiwaz* (*twz*), *I_h_ channel (ih)* and *small conductance calcium-activated potassium channel* (*SK*) (Supplementary Table 4). Main segment SCs are enriched for pattern specification process, cell-cell signaling, regionalization, and cell surface receptor signaling pathway involved in cell-cell signaling GO terms. They express a number of hormones and neuropeptide receptors (*Leucine-rich repeat-containing G protein-coupled receptor 1* (*Lgr1*), *Octα2R*, *Tachykinin-like receptor at 99D* (*TkR99D*) and *Leucokinin receptor*, (*Lkr*)), chloride channels (*SecCl* and *Chloride channel-a* (*Clc-a*)), and aquaporins (*Prip* and *Drip*) (Supplementary Table 4). The top 20 terms are displayed.

**Figure S5.**
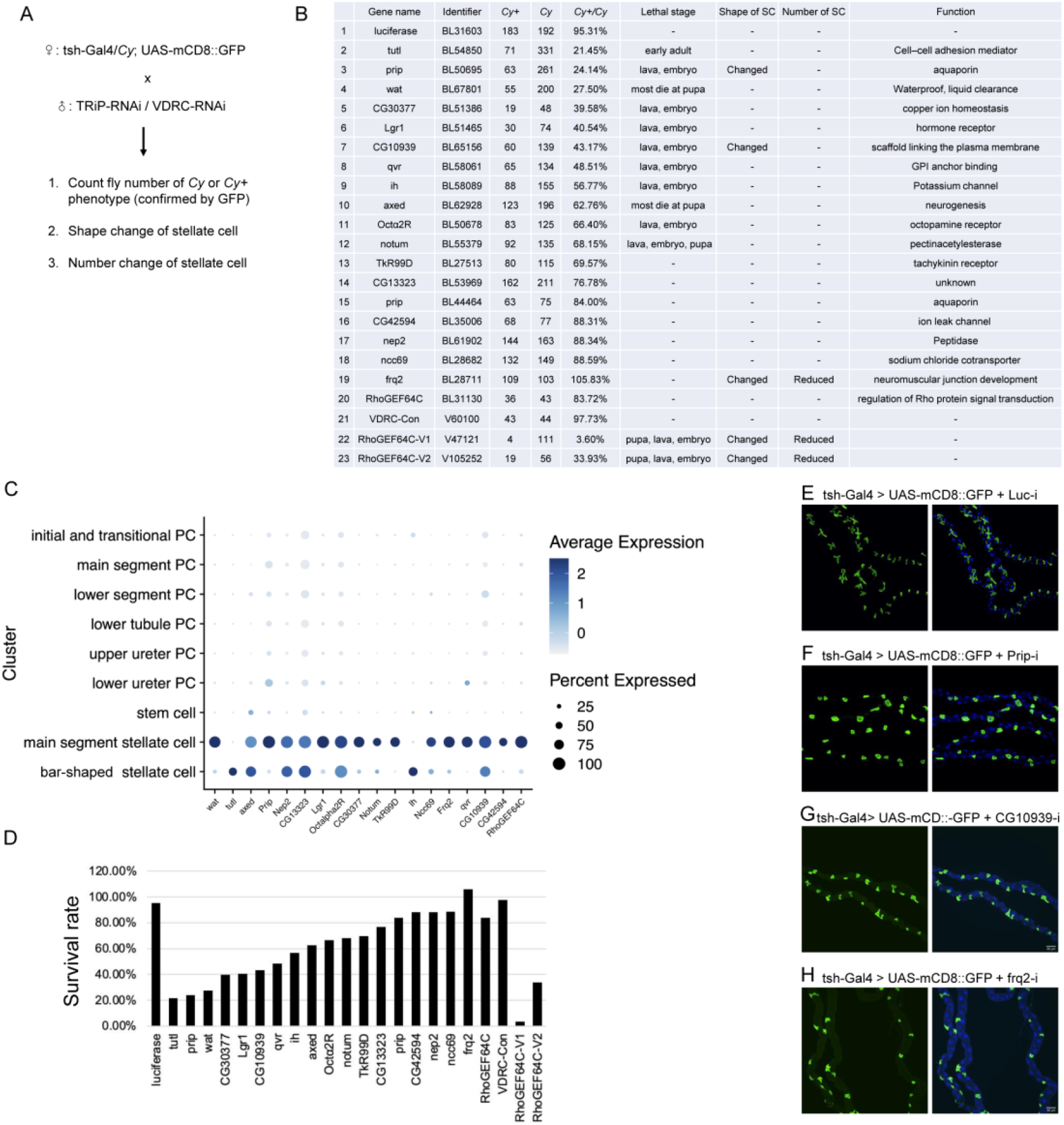
Phenotypes associate with the top stellate cell marker genes. (A) Screening strategy. SCs were visualized by tsh-Gal4 driving mCD8::GFP. (B) List of the genes tested in the screen, RNAi line identifiers, ratio of Cy+ versus Cy progenies, stage of lethality, effects on cell shape and number, and a short description of gene function. (C) Dot plots indicating the expression level of candidate genes in bar-shaped SCs and main segment SCs. (D) Histogram showing the survival rate. (E-H) SC cell shape phenotypes associated with RNAi knockdown of *Prip*, *CG10939* or *Frequenin 2* (*Frq2)*. DAPI (blue) staining for nuclei.

**Figure S6.**
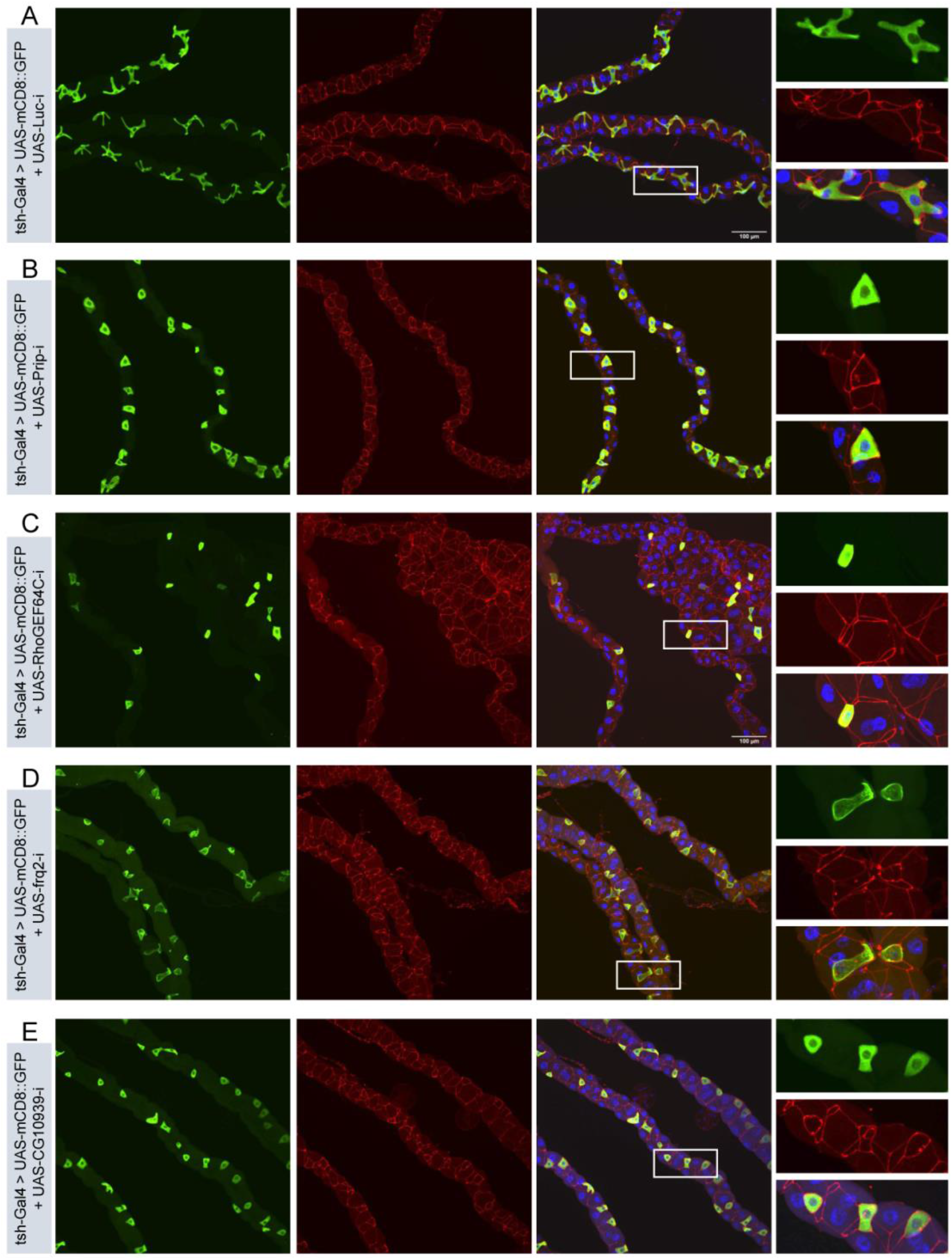
Morphology of septate junctions in Malpighian tubules. tsh-Gal4 drives mCD8::GFP expression in the adult Malpighian tubules in evenly spaced SCs. Septate junctions are labelled using anti-Dlg (red). White boxes indicate the zoomed-in regions. DAPI (blue) staining for nuclei. (A) *Luciferase* RNAi control. (B-E) Phenotypes associated with RNAi knockdown of *Prip*, *RhoGEF64c*, *Frq2* or *CG10939*. Scale bars = 100 μm.

**Figure S7.**
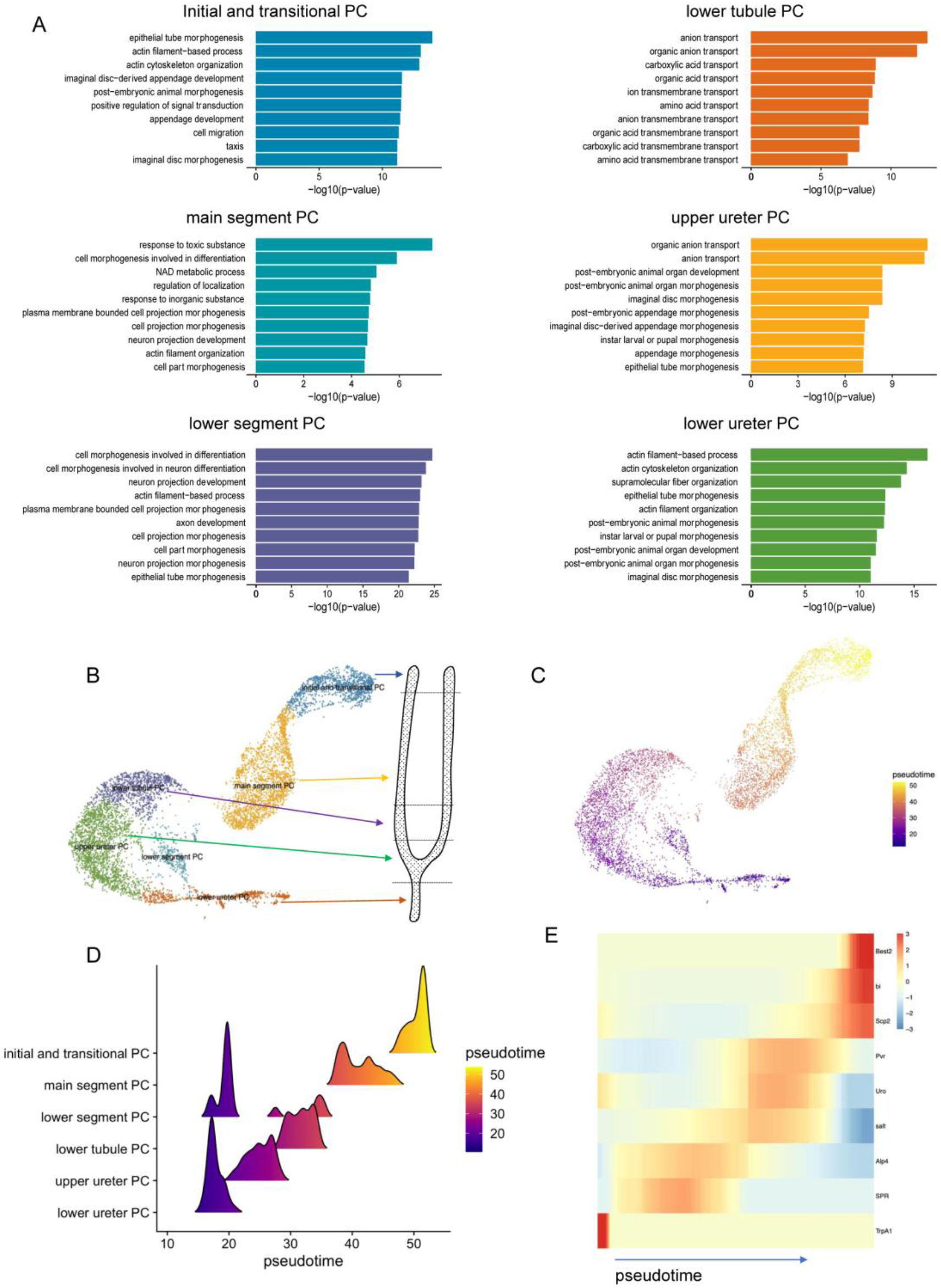
Pseudotime and GO analysis of the six PC sub-clusters. (A) UMAP of PCs showing a geographical map of the tubule. (B) Cell pseudotime was inferred using Monocle3. Purple at the beginning becomes yellow over pseudotime. (C) Sub-cell type populations for each inferred cellular trajectory. The x-axis indicates the inferred pseudotime and the y-axis indicates the height of density estimated and visualized by the RidgePlot function of Seurat R package. (D) Heatmap showing gene expression patterns during differentiation along pseudotime. (E) GO analysis of each PC cluster. Initial and transitional PCs include epithelial tube morphogenesis, actin-filament based process, and actin cytoskeleton organization. Main segment PCs include response to toxic substance, cell morphogenesis involved in differentiation, and NAD metabolic process. Lower segment PCs include cell morphogenesis involved in differentiation, cell morphogenesis involved in neuronal differentiation, and actin-filament based process GO terms. Lower tubule PCs include terms such as anion transport, organic anion transport, and carboxylic acid transport. Upper ureter PC terms include organic anion transport, and anion transport. Lower ureter PC GO terms include actin filament-based process, actin cytoskeleton organization, and supramolecular fiber organization. All the top 10 terms in lower tubule PCs were related to transport. The top 10 terms are displayed.

**Figure S8.**
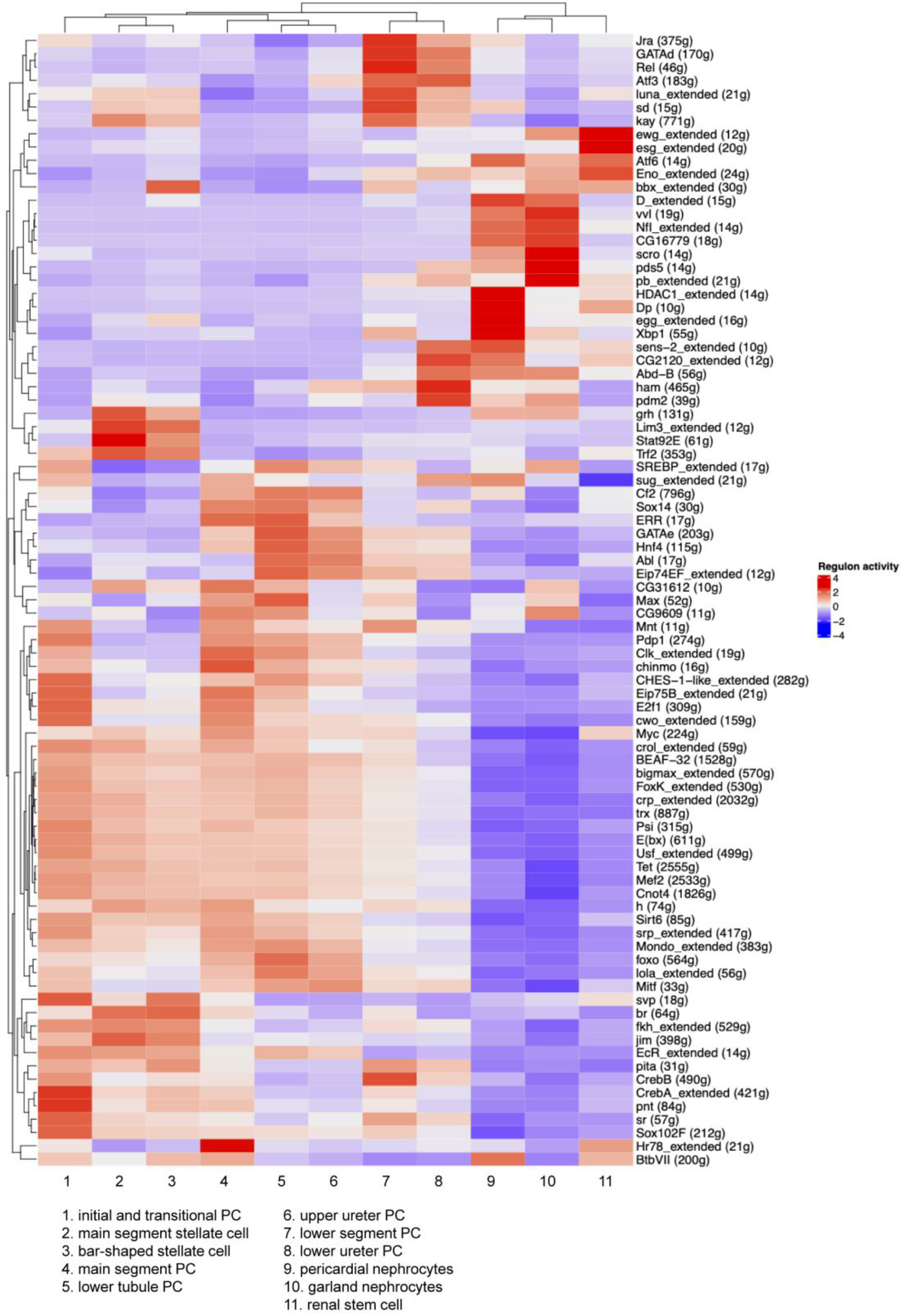
Detailed information for Fig. 4. SCENIC results for the fly kidney. Heatmap, gene expression levels in each cluster. Low regulon activity is shown in blue and high regulon activity is shown in red.

**Figure S9.**
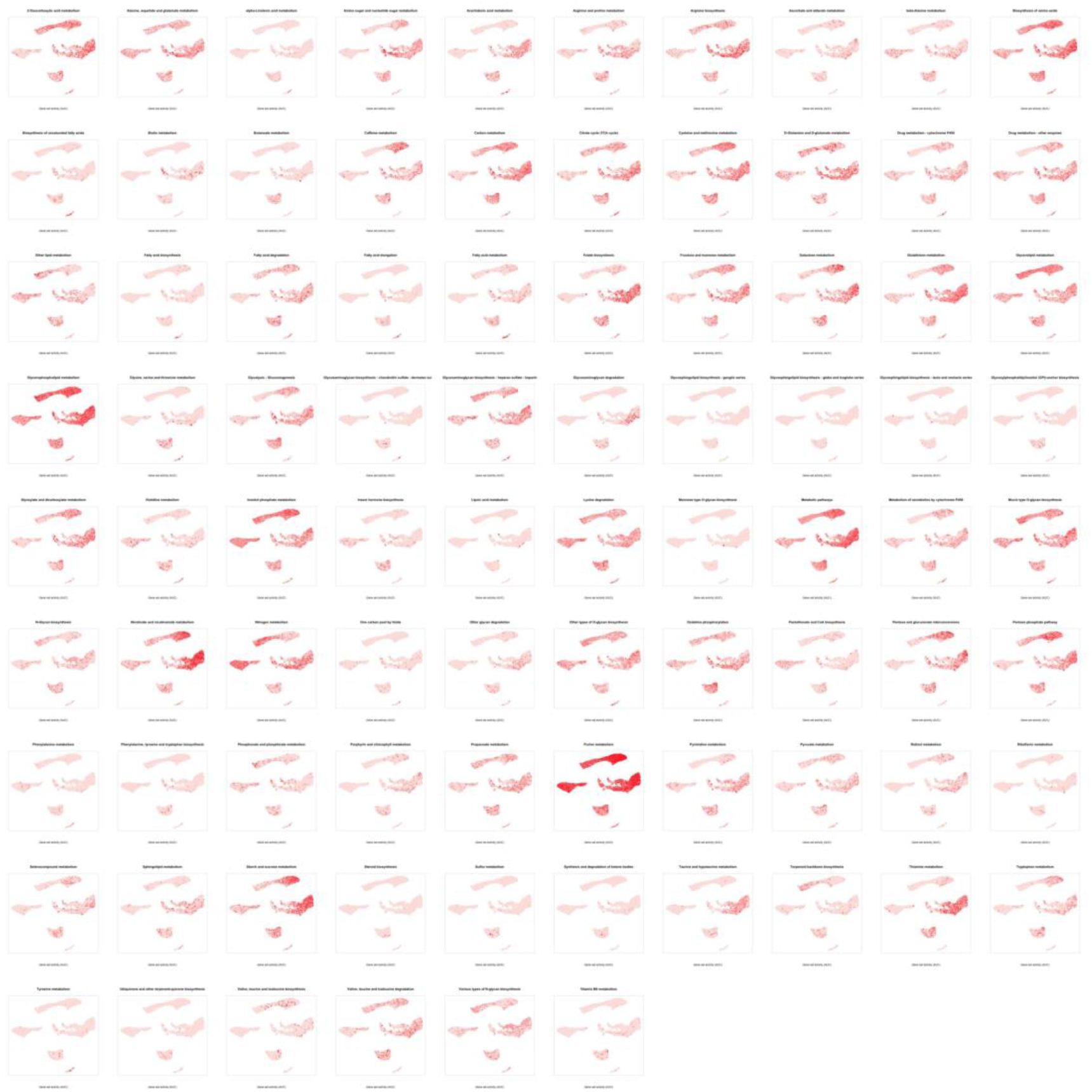
Gene set activity of 86 KEGG pathways in the UMAP fly kidney. For each pathway the color represents the gene set activity level. Red intensity reflects high gene set activity.

**Figure S10.**
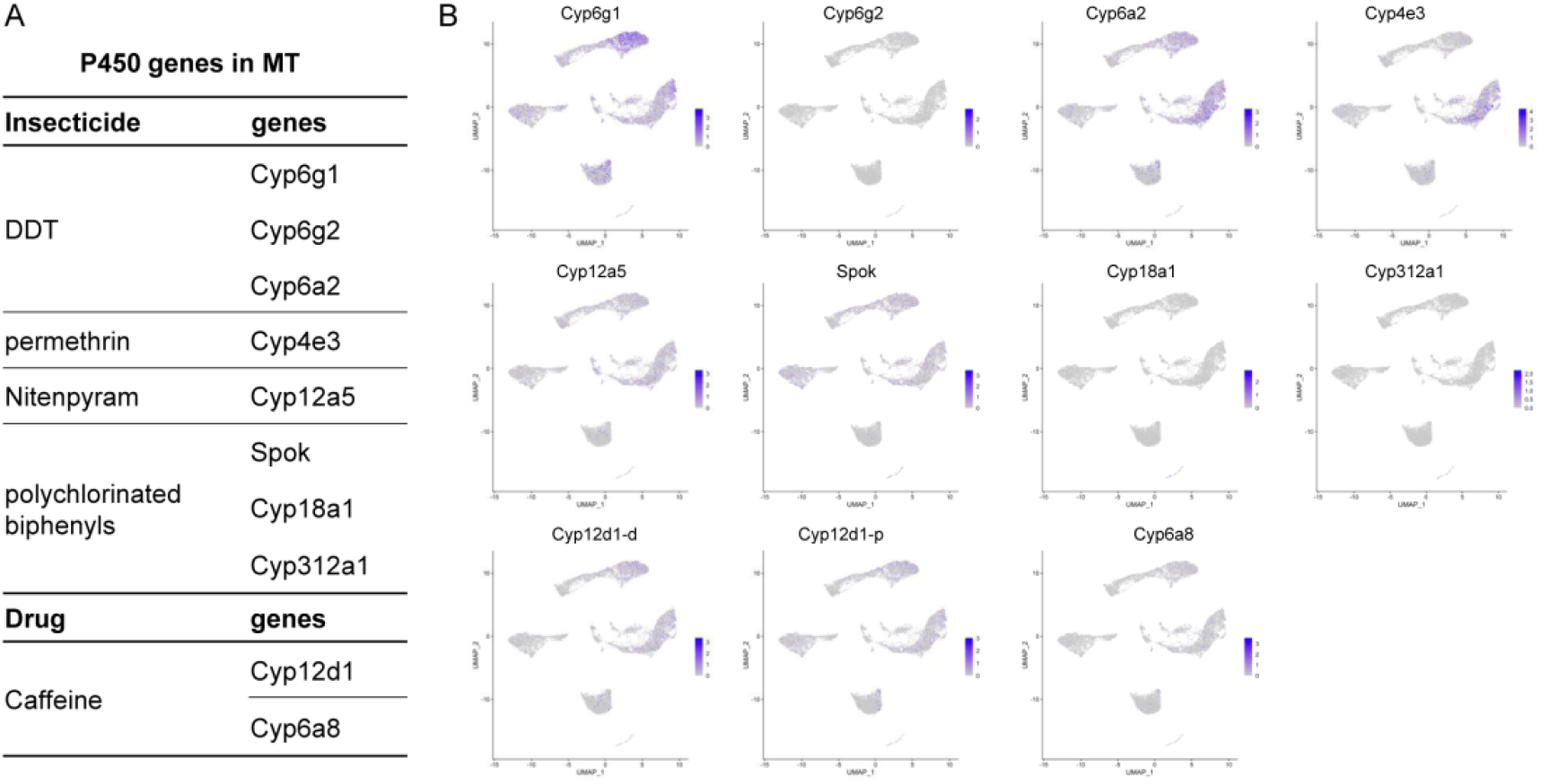
Gene expression of insecticide- and drug-related P450 genes in the fly kidney. (A) Insecticide- and drug-related P450 genes in the fly (based on Seong et al., 2020; Bergé et al., 1998; Terhzaz et al., 2015; Harrop et al. 2018; Idda et al., 2020; Najarro et al., 2015). (B) Expression levels of each P450 gene visualized by UMAP plots.

**Figure S11.**
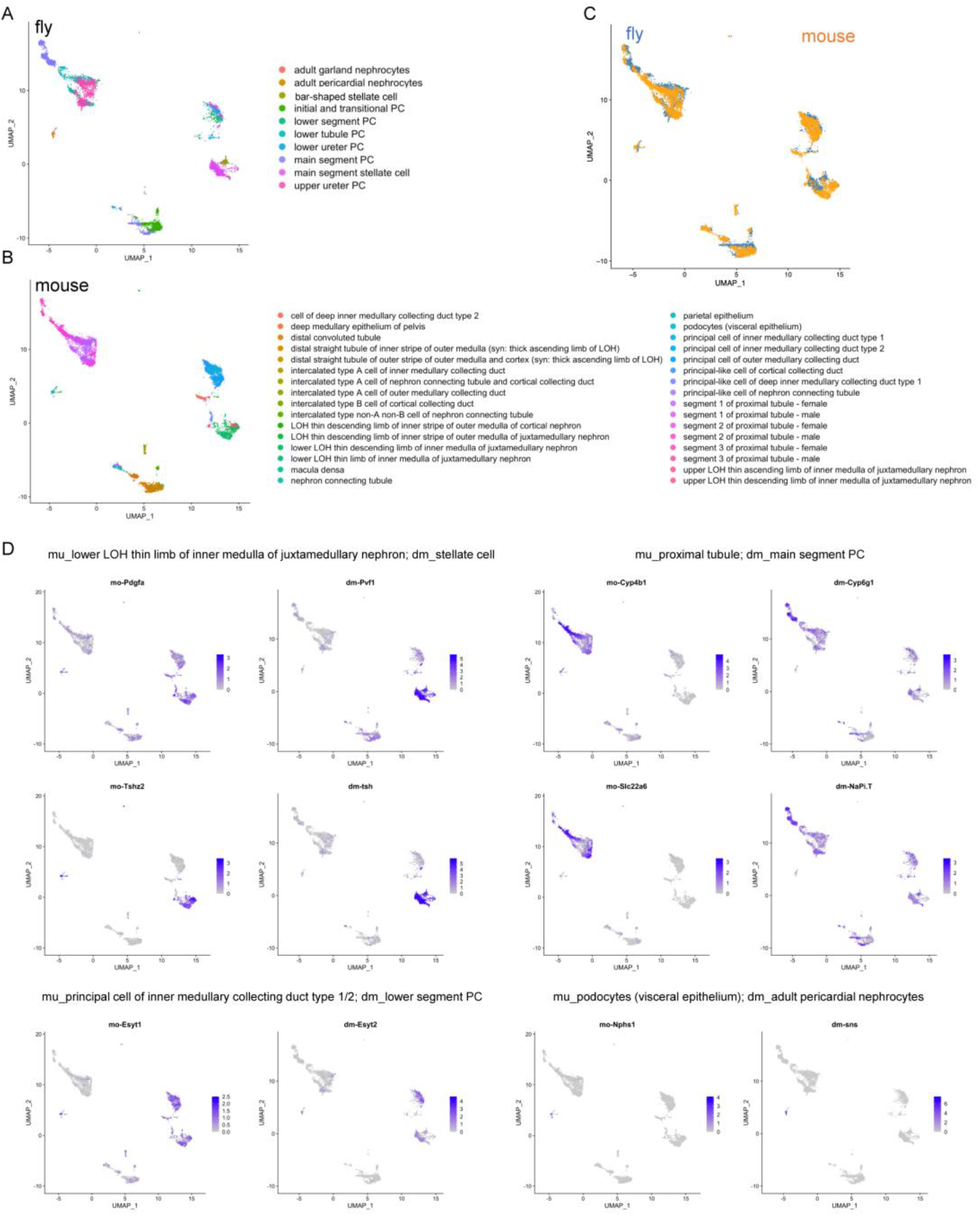
Cross-species analysis of fly kidney and mouse using SAMap. (A and B) Low dimensional representations of the cell atlases through homologous gene pairs in the mouse and fly using SAMap. (C) UMAP projection of the combined mouse (yellow) and fly (blue) manifolds. (D) Expression of orthologous gene pairs on the UMAP projection. Expressing cells are in blue and cells with no expression are shown in gray.

**Figure S12.**
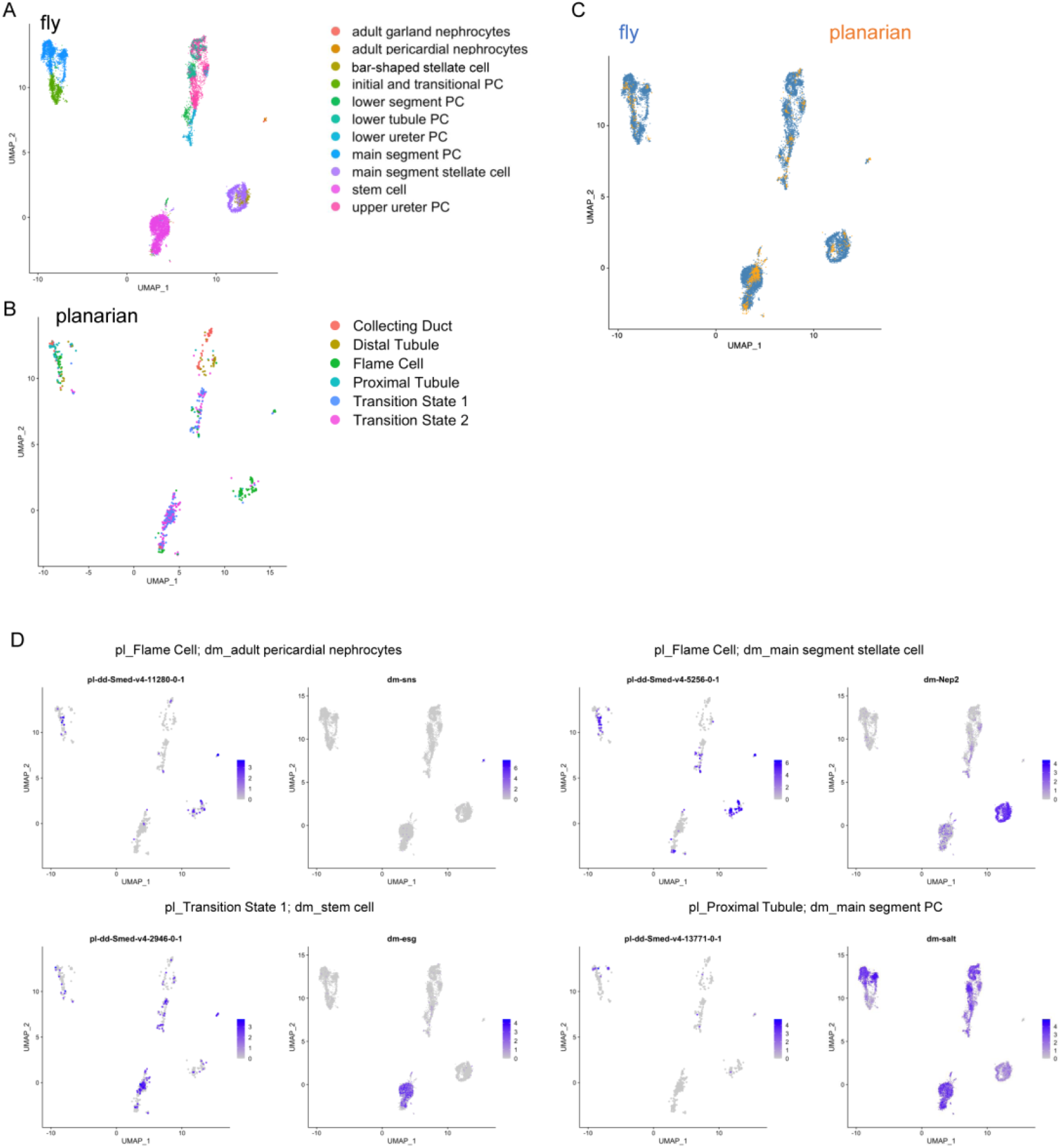
Cross-species analysis of fly kidney and planaria protonephridia using SAMap. (A and B) Low dimensional representations of the cell atlases through homologous gene pairs in the planaria and fly using SAMap. (C) UMAP projection of the combined planaria (yellow) and fly (blue) manifolds. (E) Expression of orthologous gene pairs on the UMAP projection. Expressing cells are in blue and cells with no expression are shown in gray.

**Figure S13.**
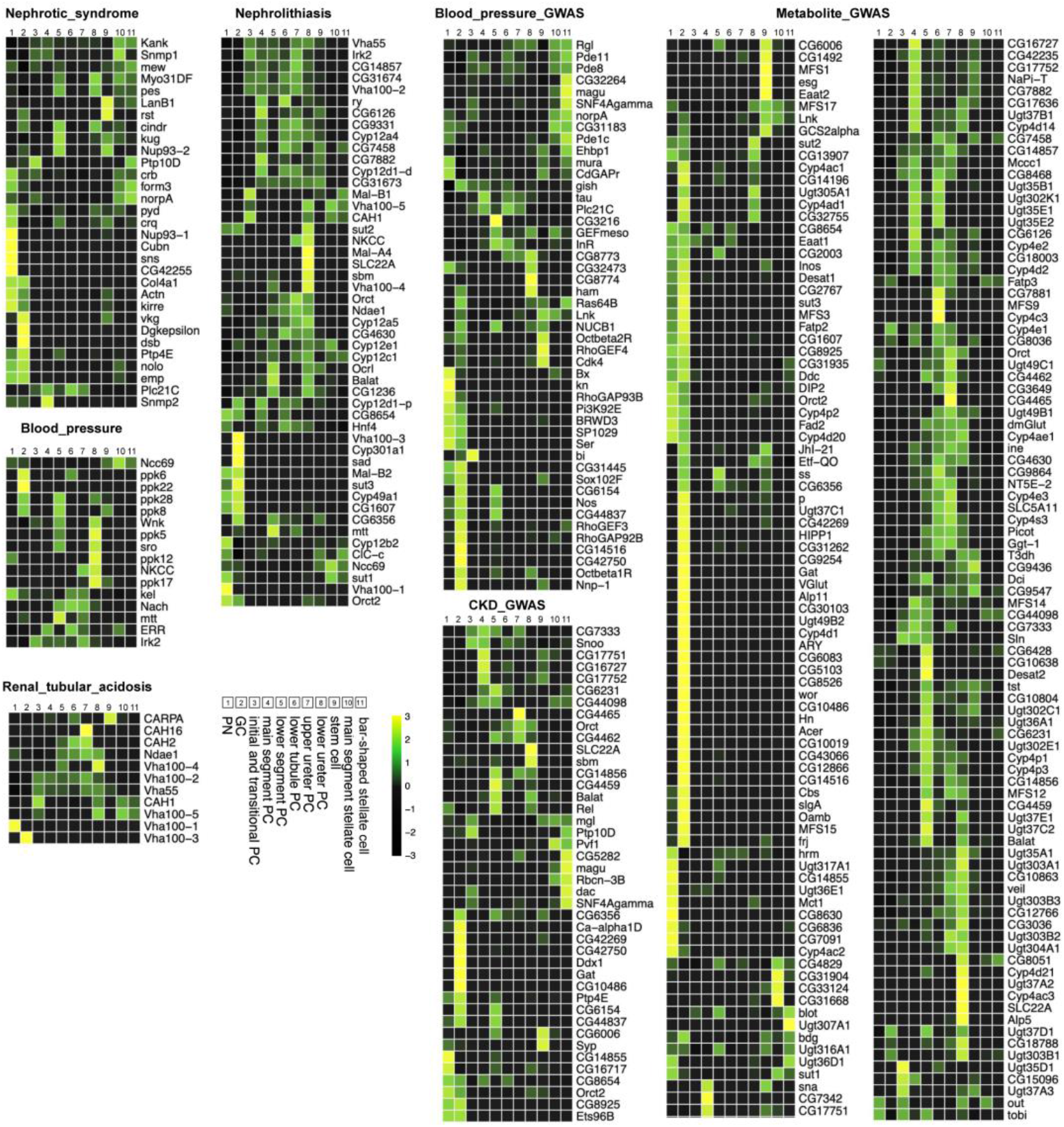
Expression of fly orthologs of human kidney disease-associated genes in specific fly kidney cell types. Average expression in single cell clusters of fly orthologs of human monogenic disease genes and complex-trait genes identified from genome-wide association studies (GWAS) (Park et al., 2018). Mean expression values of the genes were calculated for each cluster. The color scheme is based on z-score distribution (−3 < z-scores < 3). Each row in the heat map represents one gene and each column a single cell type.

**Figure S14.**
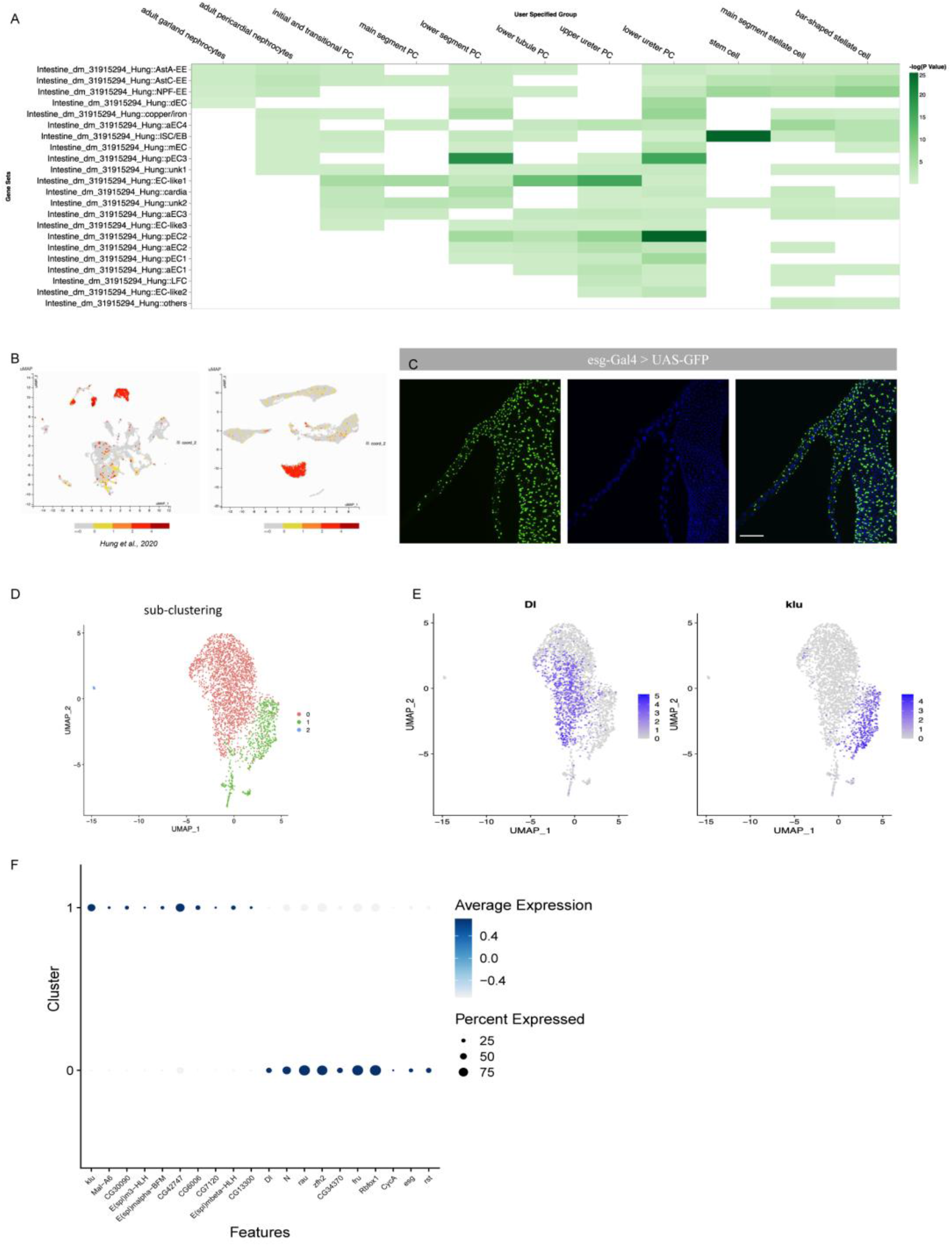
Comparison of renal stem cell and intestinal stem cell clusters. (A) snRNA-seq midgut clusters are from Hung et al. 2020. The X axis shows the 11 integrated clusters from Fig. 1B. The Y axis shows the midgut clusters from Hung et al. (2020). Colors represent gene expression similarities. Note that RSCs are highly similar to ISCs. In addition, lower ureter PCs share high similarity with pEC2 (posterior enterocytes). (B) *escargot (esg)* expression in the gut and Malpighian tubule UMAPs. (C) *esg* expression in the Malpighian tubules visualized using esg-Gal4 driving UAS-GFP expression. Scale bars = 100 μm. (D) UMAP distribution of different sub-clusters of RSCs. (E) *Delta (Dl)* and *klumpfuss (klu)* expression in the RSC UMAP subclusters. (F) Dot plot showing the expression levels and percentage of cells expressing the various markers in the *Dl* and *klu* sub-clusters.

**Figure S15.**
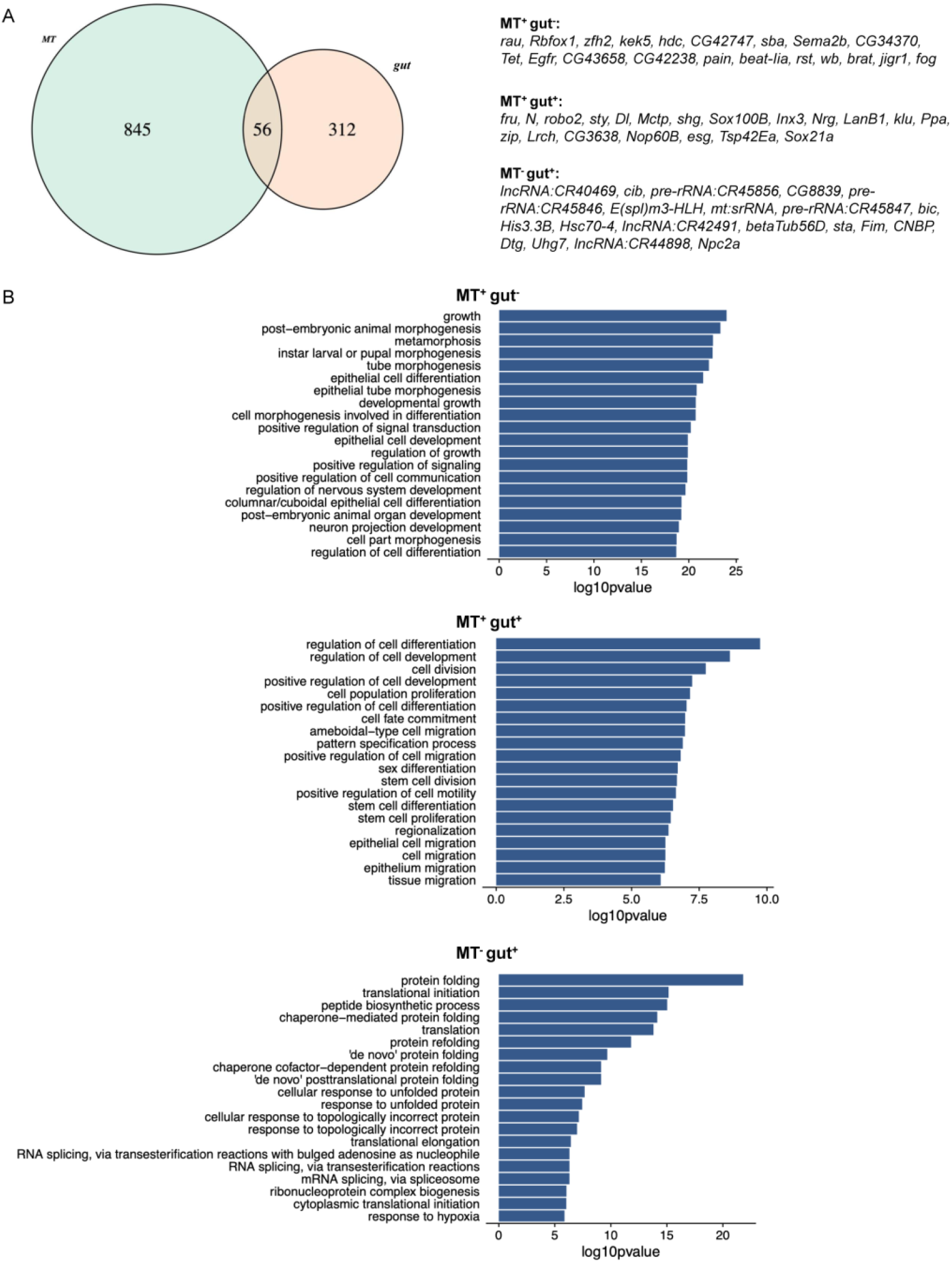
Comparison of renal stem cell and intestinal stem cell clusters. (A) Venn diagram of the overlap between Malpighian tubule (MT) and gut stem cell top marker genes (log2FC > 0.25 and adjust p-value < 0.05). On the right are identities of the top genes in the three regions of the diagram. (B) Gene Ontology (GO) analysis of MT^+^gut^-^, MT^+^gut^+^ and MT^-^gut^+^. The top 20 terms are displayed.

**Figure S16.**
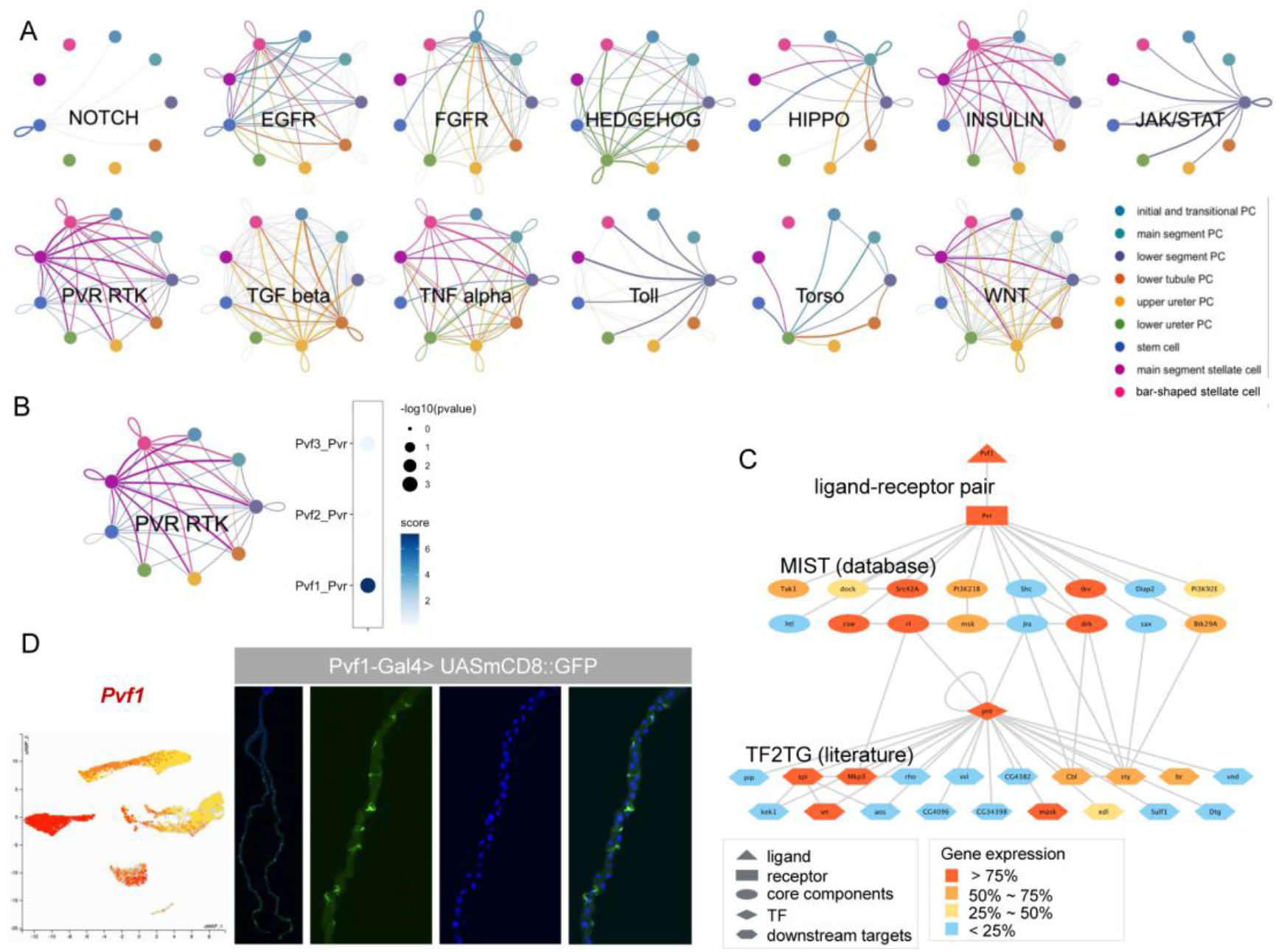
Cell–cell communication analysis in the adult fly kidney. (A) Network of 13 signaling pathways in fly kidney cell clusters. Each of the 11 cell clusters is displayed in a different color. The predicted interaction between two clusters is indicated by a color curve. The thickness of the curve indicates the strength of the interaction. The full list of predicted ligand/receptor pair genes can be found in Supplementary Table 12. (B) Ligand-receptor interaction between *Pvf1* and its receptor *Pvr* in main segment SCs and main segment PCs. The panel on the right shows a dot plot of the interaction score and specificity of ligand-receptor pairs between main segment principal cell cluster and main segment stellate cell cluster. (C) *Pvr*-*pvf1* interaction network based on MIST and TF2TG. (D) *pvf1* expression in SCs visualized using pvf1-Gal4 driving UAS-mCD8::GFP expression.

**Supplementary Table legends:**

**Supplementary Table 1.** Basic statistics of snRNA-seq libraries.

**Supplementary Table 2.** Differentially expressed genes in each cluster. Only positive marker genes are shown.

**Supplementary Table 3.** Table of validated markers, from previous studies and this study, allowing assignment of clusters to cell types or regions.

**Supplementary Table 4.** GO terms comparison of bar-shaped SCs and main segment SCs.

**Supplementary Table 5.** GO terms of the six PC cell clusters.

**Supplementary Table 6.** Full list of regulons and their respective predicted target genes.

**Supplementary Table 7.** List of cell type-specific transcription factors.

**Supplementary Table 8.** Gene pairs for fly and mouse cell type mappings.

**Supplementary Table 9.** Gene pairs for fly and planarian cell type mappings.

**Supplementary Table 10.** Gene list comparison of RSCs and ISCs.

**Supplementary Table 11.** GO terms comparison of RSCs and ISCs.

**Supplementary Table 12.** List of gene pairs for cell-cell communication predictions.

